# Androgen Receptor mediates beige adipocyte homeostasis and plasticity

**DOI:** 10.64898/2026.09.16.750644

**Authors:** Rajesh Sahu, Qingshuang Cai, Manon Ohlmann, Pierrick Dupre, Quentin Brassart, Anthony Bringolf, Joe G. Rizk, Anouk Charlot, Tao Ye, Sirine Souali-Crespo, Kamar C. Ghaibour, Eric Metzger, Roland Schüle, Matthieu Lacroix, Laurent Le Cam, Joffrey Zoll, Daniel Metzger, Delphine Duteil

## Abstract

Male adipose tissue undergoes profound post-pubertal remodeling characterized by the progressive transition of beige adipocytes toward a white adipocyte phenotype. Although androgen signaling has been implicated in adipose tissue biology, its role in beige adipocyte remodeling and metabolic plasticity remains poorly understood. Here we show that androgen receptor (AR) signaling is dynamically activated in inguinal white adipose tissue during the post-pubertal beige-to-white transition in male mice. Inducible deletion of AR in beige adipocytes impaired this remodeling process, resulting in the persistence of multilocular beige-like adipocytes despite reduced thermogenic competence and marked mitochondrial abnormalities. Transcriptomic and cistromic analyses identified AR as a direct regulator of adipocyte metabolic and differentiation programs, whereas immune-related transcriptional signatures in AR-deficient adipose tissue primarily reflected macrophage infiltration and inflammatory remodeling. Loss of AR promoted mitochondrial dysfunction, mitophagy, and altered glucose handling, while cell-autonomous AR silencing in beige adipocytes recapitulated key defects in mitochondrial organization and adipocyte identity. Longitudinal and metabolic challenge studies further demonstrated that AR signaling is required for age-associated adipose remodeling and adaptive beige adipocyte plasticity during high-fat diet feeding and cold exposure. Together, these findings identify AR as a central regulator of beige adipocyte remodeling, mitochondrial homeostasis, and adaptive metabolic function in male adipose tissue.

## INTRODUCTION

Adipose tissue is a central organ for energy homeostasis and endocrine regulation. Based on the characteristics of resident adipocytes, it is broadly classified into brown adipose tissue (BAT) and white adipose tissue (WAT). BAT, located predominantly in the interscapular region in mice [1], is composed of brown adipocytes specialized for thermogenesis through lipid oxidation. These cells contain multilocular lipid droplets and are highly enriched in mitochondria, reflecting their high metabolic activity. In contrast, WAT primarily functions in energy storage, thermal insulation, adipokine secretion, and is composed of adipocytes containing a large unilocular lipid droplet and relatively few mitochondria. Specific WAT depots such as the inguinal WAT (ingWAT) consist of a third adipocyte subtype, termed beige adipocytes. Despite residing within WAT, beige adipocytes closely resemble brown adipocytes both morphologically and functionally. Although many studies have focused on inducible beige adipocytes formed in response to environmental stimuli, mice also possess a population of basal beige adipocytes under unstimulated conditions [2]. Beige adipocytes can also transdifferentiate from mature white adipocytes into brown-like adipocytes following cold exposure [3], or administration of the β3-adrenergic receptor agonist CL-316243 [4]. Importantly, transdifferentiated beige adipocytes retain considerable plasticity, as they can revert to a white adipocyte-like morphology and transcriptional state at ambient temperature and subsequently re-acquire beige characteristics upon repeated cold exposure [5]. Conversely, aging promotes beige-to-white transdifferentiation [6], accompanied by increased lipid accumulation, impaired energy utilization [7], and a diminished capacity to induce beige adipogenesis in response to cold exposure [8]. The homeostasis in adipose tissue is maintained via both adipocyte intrinsic gene programs, as well as external signals from its microenvironment, which consists of various non-adipocyte cell types such as immune cells and progenitors, in addition to the ECM and vasculature.

Besides testes and adrenal glands, adipose tissue is also an important organ for androgen synthesis [9], with the most abundant one being dehydroepiandrosterone (DHEA) instead of testosterone [10]. Androgens modulate adipocyte physiology in a depot-specific manner in males; testosterone was found to inhibit fatty acid uptake [11] and lipoprotein lipase (LPL) activity [12], while increasing lipolysis [13] in specific human and mice fat depots. Studies have also shown correlation between circulating androgen levels and adiposity, where obese men exhibit a decrease in testosterone levels [14]. Androgens’ physiological effects are mediated via the androgen receptor (AR) in adipocytes [15], a ligand-dependent nuclear receptor located in the cytoplasm in the absence of hormone. Upon androgen binding, AR gets activated, undergoes structural changes, translocate into the nucleus, and binds to specific DNA sequences termed androgen response elements (AREs) to modulate gene transcription [16-18].

Given the remarkable plasticity of beige adipocytes, the regulatory mechanisms governing their maintenance and post-pubertal transdifferentiation remain incompletely understood. More importantly, these regulatory mechanisms have not been studied from the perspective of AR. In this study, we combine histological, transcriptomic, and metabolic analyses to identify AR as a key regulator of beige adipocyte maintenance and plasticity, nutrient metabolism, and immune remodeling within ingWAT.

## RESULTS

### AR promotes post-puberty beige adipocyte whitening in male mice

To investigate the role of androgen receptor (AR) signaling in adipocytes during post-pubertal adipose tissue remodeling in male mice, we first characterized AR expression across adipose depots during the period of beige-to-white transition. To this aim, we performed immunohistochemical detection of AR levels in mouse inguinal white adipose tissue (ingWAT), epididymal white adipose tissue (epWAT), and brown adipose tissue (BAT) from 8-to 15-weeks (Fig. 1A, 1B and Fig. S1A). In ingWAT, while cytoplasmic AR levels remained relatively stable over time, the nuclear AR fraction markedly increased at 9 weeks of age, followed by a decline at 10 and 15 weeks of age, coinciding with the physiological beige-to-white adipocyte transition (Fig. 1A and 1B). In contrast, nuclear AR expression remained negligible in BAT and relatively stable in epWAT across ages (Fig. S1A), revealing depot-specific and age-dependent AR localization patterns. RT-qPCR and western blot analyses at 10 weeks of age further confirmed depot-specific differences in AR expression (Fig. S1B and S1C), in agreement with previous reports [19].

**Figure 1:**
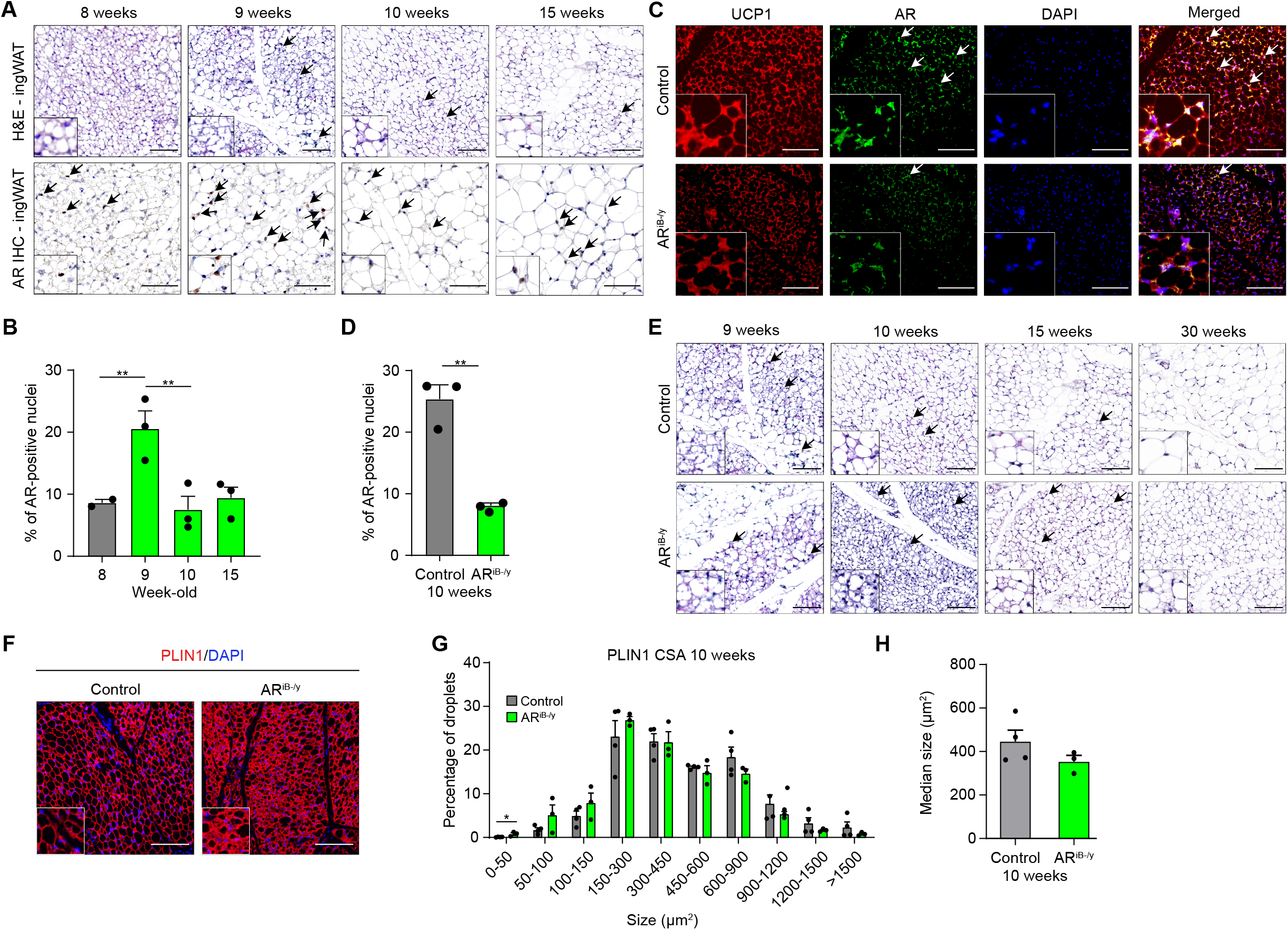
AR promotes post-puberty beige adipocyte whitening in male mice. (A-B) Hematoxylin and Eosin (H&E) staining and immunohistochemical detection of the androgen receptor (AR) in the inguinal white adipose tissue (ingWAT) of control mice at 8, 9, 10, and 15 weeks of age (A), and corresponding quantification of the percentage of AR-positive nuclei (B). Insets at the lower-left corner represent higher magnification of corresponding images. (A) Scale bars, 1001µm. Black arrows indicate dense lipid droplet clusters, representative of beige adipocytes. (B) Black arrows point to AR expression in the nucleus. Scale bars, 50⍰µm. Data are represented as mean + SEM. Statistical test used is ordinary one-way ANOVA. **, p < 0.01. (C-D) Immunofluorescent detection of uncoupling protein 1 (UCP1; red) and AR (green) in ingWAT of 10-week-old control and AR^iB-/y^ mice (C), and corresponding quantification of the percentage of AR-positive nuclei (D). Nuclei were stained with DAPI. White arrows indicate AR nuclear expression. Insets at the lower-left corner represent higher magnification of corresponding images. Scale bars, 100⍰µm. Data are represented as mean + SEM. Statistical test used is two-tailed unpaired t-test. **, p < 0.01. (E) Hematoxylin and Eosin (H&E) staining of ingWAT at 9 and 10 weeks of age for control and AR^iB-/y^ mice. Black arrows indicate dense lipid droplet clusters, representative of beige adipocytes. Insets at the lower-left corner represent higher magnification of corresponding images. Scale bars, 1001µm. (F-H) Immunofluorescent detection of Perilipin (PLIN1; red) (F), and corresponding quantification of lipid droplet cross-sectional area (CSA) (G) and median size (H), in ingWAT of 10-week-old control and AR^iB-/y^ mice. Nuclei were stained with DAPI. Insets at the lower-left corner represent higher magnification of corresponding images. Scale bars, 1001µm. Data are represented as mean + SEM. Statistical test used are Two-way ANOVA with Tukey post-hoc correction (G) and two-tailed unpaired t-test (H). *, p < 0.05.

Given the temporal enrichment of nuclear AR in ingWAT during the early stage of beige adipocyte remodeling, we hypothesized that AR activation may contribute to the acquisition of a mature adipocyte phenotype during the beige-to-white transition. To test this hypothesis, we intercrossed AR-floxed animals with *Ucp1*-CreER^T2^ mice to generate AR^iB-/y^ mice. CreER^T2^-mediated recombination was induced by intraperitoneal tamoxifen administration at 9 weeks to selectively delete AR in UCP1-expressing brown and beige adipocytes (Fig. S1D). Efficient recombination of *Ar* exon 1 was confirmed by genotyping PCR (Fig. S1E), and AR/UCP1 co-staining by immunofluorescence (IF) demonstrated selective loss of AR expression in UCP1-positive adipocytes at 10 weeks of age, one week after AR ablation (Fig. 1C and 1D).

Histological analysis revealed that ingWAT morphology from 9-week-old pre-mutant mice was comparable to that of control littermates prior to tamoxifen injection (Fig. 1E). As expected, control mice underwent the characteristic age-associated beige-to-white transition between 9 and 10 weeks, with progressive acquisition of large unilocular lipid droplets. In contrast, AR-deficient ingWAT retained abundant multilocular adipocytes at 10 weeks, indicating that AR deletion prevents the whitening process (Fig. 1E). Importantly, this phenotype persisted beyond the initial transition period, as multilocular adipocytes remained detectable at later ages, whereas ingWAT from control littermates progressively acquired a white adipocyte morphology (Fig. 1E). AR deletion in beige and brown adipocytes had no detectable effect on epWAT morphology, whereas BAT displayed reduced lipid droplet sizes relative to control animals (Fig. S1F and S1G). This remodeling defect was not associated with major alterations in adipose depot mass, as adipose tissue weights remained comparable between genotypes at the analyzed time points (Fig. S1H and S1I). Smaller lipid droplets in BAT could thus be a result of altered lipid accumulation/turnover rather than altered cellular abundance in tissue. Consistent with the persistence of beige-like adipocytes in ingWAT, PLIN1 staining combined with lipid droplet morphometry demonstrated a significant enrichment of small lipid droplets in ARiB-/y mice compared with control littermates (Fig. 1F-H). Together, these findings demonstrate that AR signaling is required for functional remodeling of beige adipocytes during post-pubertal ingWAT maturation.

### AR directly controls adipocyte metabolic programs driving beige adipocyte remodeling

Having established that AR is required for physiological beige adipocyte whitening, we next sought to define the transcriptional mechanisms underlying this phenotype. As AR functions as a ligand-activated transcription factor, we first characterized its genomic occupancy in ingWAT from control mice at 10 weeks of age. AR cistrome characterization by ChIP-seq identified 9,287 AR-binding sites (ARBS), associated with 5,265 annotated genes (Fig. S2A). AR occupancy was predominantly detected at distal genomic regions, including intergenic (67%) and intronic (31%) regions, whereas only 2% of AR peaks localized to proximal promoter regions [transcription start site (TSS), −1 kb to +100 bp], consistent with preferential enhancer binding by AR (Fig. S2A and 2B). Integration with H3K4me2 ChIP-seq revealed that the majority of AR-bound genes were associated with active chromatin regions, supporting a direct role of AR in regulating transcriptional programs in beige adipocytes (Fig. 2A and 2B). Motif enrichment analysis revealed that 64% and 35% of AR peaks contained canonical AR half-site and full-site motifs, respectively, confirming the specificity of AR recruitment to DNA (Fig. 2C). Pathway analysis of genes associated with ARBS revealed enrichment for pathways related to cell differentiation and homeostasis (Fig. 2D), as exemplified by Sorbs1, Gcg, Creb5, Adcy1, Ins1, and Glp1r (Fig. 2E and S2C).

**Figure 2:**
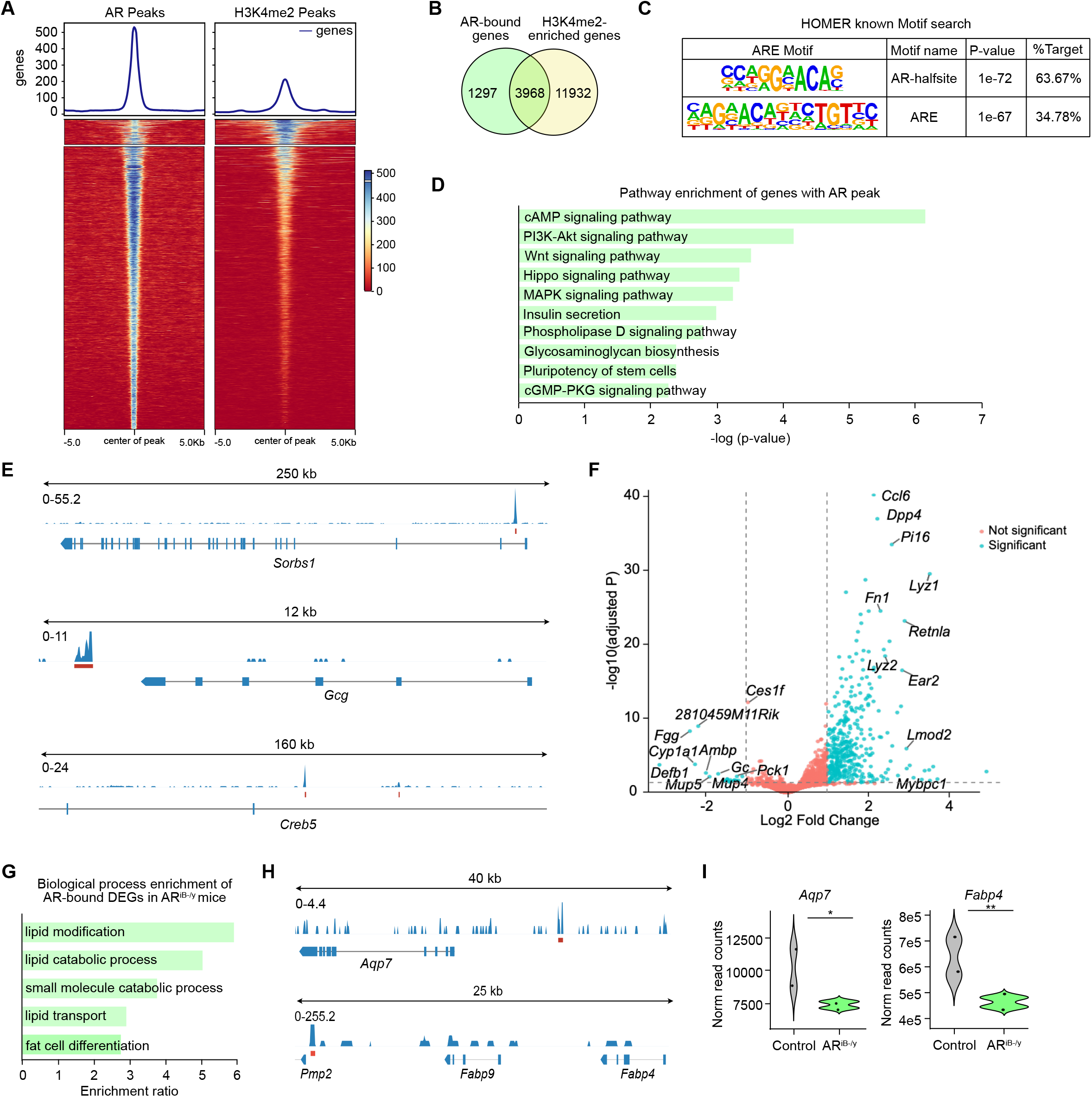
AR directly controls adipocyte metabolic programs driving beige adipocyte remodeling. (A) Tag density map of AR and H3K4me2 in ingWAT, +/-5 kb from the AR peak center sorted by H3K4me2 peak length, and corresponding average tag density profiles. (B) Overlap between AR-bound and H3K4me2-enriched genes in ingWAT of 10-week-old AR^iB-/y^ mice. (C) HOMER motif analysis of AR binding sites. p-value: hypergeometric testing. (D) Pathway analysis of AR-bound genes in ingWAT of 10-week-old control mice. (E) AR localization at representative loci on the chromatin of ingWAT by ChIP-seq. (F) Volcano plot depicting in cyan the genes differentially expressed between AR^iB-/y^ and control ingWAT at 10 weeks. Selected markers are annotated. Additional genes are indicated in red. (G) Gene ontology analysis for biological processes enriched from genes differentially expressed in the ingWAT between control and AR^iB-/y^ mice with an AR peak within a maximum of 100 kbp distance from their TSS. (H-I) AR localization at *Aqp7* and *Fabp4* in ingWAT (H), and corresponding violin plots for normalized read counts from RNA-seq data of ingWAT in 10-week-old control and AR^iB-/y^ mice (I). P-values were obtained from DESeq2. *, p < 0.05; **, p < 0.01.

To determine which AR-associated genes were transcriptionally affected by AR loss, we next integrated AR ChIP-seq and RNA-seq datasets from control and AR^iB-/y^ ingWAT at 10 weeks of age. Transcriptomic analysis identified 2,150 differentially expressed genes (DEGs) following AR deletion, including 1,477 upregulated and 673 downregulated genes (Fig. 2F). Because AR predominantly binds distal enhancer regions rather than promoters, we opted for a gene-centric integration strategy to identify putative direct AR targets among these differentially expressed genes. ARBS located within a 100 kb window of transcriptionally altered genes were considered associated with potential AR regulation. Using this approach, 42.8% of DEGs were associated with at least one AR-binding site (Fig. S2D), indicating that loss of AR affects genes directly controlled by AR. Gene ontology analysis of these putative direct AR targets revealed enrichment for pathways involved in lipid metabolism and adipocyte differentiation (Fig. 2G). Notably, several genes identified from the AR cistrome, including Fabp4, Aqp7, Apoc3, and *Adrb3*, were both associated with AR-binding sites and transcriptionally reduced following AR deletion (Fig. 2H, 2I and S2E).

Consistent with these transcriptional alterations, pathway analysis of the complete RNA-seq dataset revealed suppression of metabolic pathways following AR ablation, including lipid metabolism, PPAR signaling, glutathione metabolism, and peroxisomal pathways, while up-regulated genes were mainly associated with immune reaction signature (Fig. S2F and S2G). To further identify transcriptional regulators associated with AR-dependent gene expression changes, we performed LISA transcription factor prediction analysis on the differentially expressed genes identified in AR^iB-/y^ ingWAT. Downregulated genes were preferentially associated with AR and PPARG as the predominant predicted regulators of downregulated genes. In contrast, upregulated genes were associated with transcriptional regulators including glucocorticoid receptor (GR) and macrophage-associated factors such as IRF4 and IRF8, suggesting the emergence of compensatory transcriptional programs following AR loss (Fig. S2H). Together, our data demonstrate that AR directly regulates a transcriptional network required for maintenance of beige adipocyte identity and lipid metabolic programs, while loss of AR triggers broader compensatory transcriptional remodeling within the ingWAT microenvironment.

### AR deficiency disrupts mitochondrial metabolic programs and induces mitochondrial stress in beige adipocytes

Given that AR-dependent transcriptional programs were enriched for metabolic pathways (Fig. 2G), we next investigated the consequences of AR loss on the mitochondria in beige adipocytes. Transcriptomic analysis revealed a broad remodeling of mitochondrial and metabolic programs in AR^iB-/y^ ingWAT. Indeed, genes involved in mitochondrial substrate utilization and oxidative metabolism were reduced following AR deletion, including β-oxidation markers *Acads, Acat1, Acox1, Acot12, Cpt2, Decr1*, and *Etfa* (Fig 3A). In parallel, transcripts associated with mitochondrial respiratory function and electron transport chain activity were decreased, including *Ndufab1, Ndufb3, Ndufv3*, from complex I, and coenzyme Q related factors *Coq4* and *Coq8b*, indicating altered mitochondrial bioenergetic capacity.

**Figure 3:**
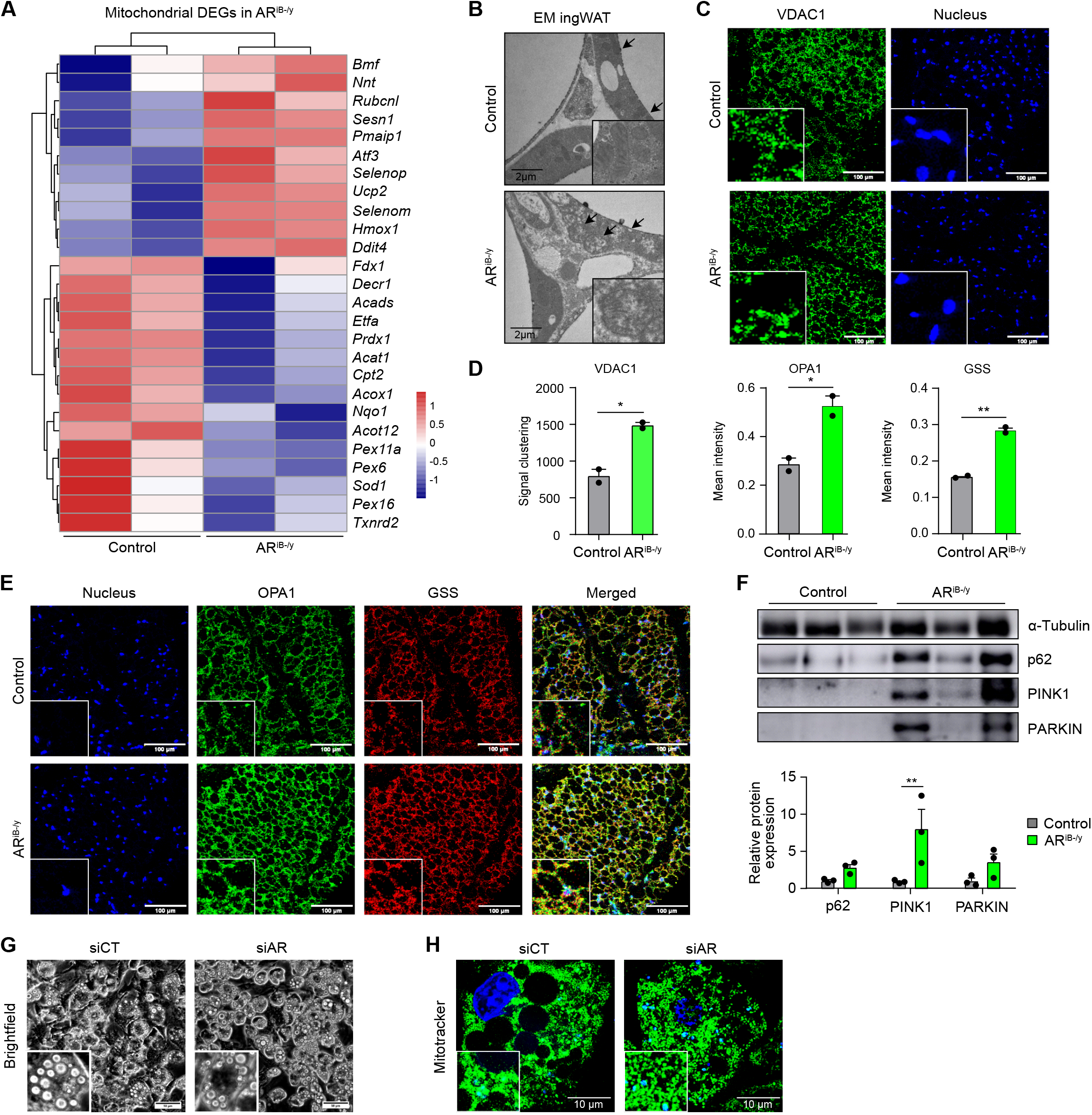
AR deficiency disrupts mitochondrial metabolic programs and induces mitochondrial stress in beige adipocytes. (A)Heatmap depicting the mean-centered normalized expression of mitochondria-associated genes obtained from the RNA-seq datasets of ingWAT from control and AR^iB-/y^ mice at 10 weeks of age. (B)Ultrastructure analysis of ingWAT from 10-week-old control and AR^iB-/y^ mice. Black arrows indicate distinct mitochondria. Insets at the lower-right corner represent higher magnification of corresponding images. Scale bars, 1⍰µm. (C-D) Imaging mass cytometry detection of VDAC1 (green) (C) and corresponding quantification of signal clustering (GLCM contrast algorithm) (D) in ingWAT of 10-week-old control and AR^iB-/y^ mice. Nuclei are visible in blue. Insets at the lower-left corner represent higher magnification of corresponding images. Scale bars, 100⍰µm. Data are presented as mean + SEM. Statistical test used is two-tailed unpaired t-test. *, p < 0.05; **, p < 0.01. (E)Imaging mass cytometry detection of OPA1 (green) and Glutathione synthase (GS, red), and corresponding quantification (D) of mean fluorescence intensity, in ingWAT of 10-week-old control and AR^iB-/y^ mice. Nuclei are visible in blue. Insets at the lower-left corner represent higher magnification of corresponding images. Scale bars, 100⍰µm. Data are presented as mean + SEM. Statistical test used is two-tailed unpaired t-test. *, p < 0.05; **, p < 0.01. (F)Western blot analysis and corresponding quantification of indicated proteins in ingWAT of 10-week-old control and AR^iB-/y^ mice. Tubulin was used as a loading control. Data are presented as mean + SEM. Statistical test used is two-tailed unpaired t-test. **, p < 0.01. (G-H) Brightfield images (G) and mitotracker staining (H) of control-(siCT) and AR-siRNA transfected (siAR) immortalized beige adipocytes, 3-days post-differentiation. Insets at the lower-left corner represent higher magnification of corresponding images. Nuclei (blue) are stained with Hoechst. Scale bars, 50⍰µm (G) and 10⍰µm (H).

AR deficiency also affected pathways involved in peroxisomal lipid metabolism and redox homeostasis, as demonstrated by reduced expression of peroxisome assembly (e.g *Pex11a, Pex16, Pex6*) and antioxidant-associated genes (e.g. *Txnrd2, Prdx1, Sod1*, and *Nqo1*) (Fig. 3A) Conversely, AR-deficient ingWAT displayed increased expression of several mitochondrial stress and metabolic adaptation genes, including *Ddit4, Sesn1, Atf3, Hmox1, Ucp2, Nnt, Selenop*, and *Selenom* (Fig. 3A). These genes are associated with cellular responses to energetic stress, redox imbalance, and activation of compensatory pathways, indicating that AR loss triggers an adaptive response to mitochondrial dysfunction rather than a simple loss of mitochondrial activity. Finally, genes associated with mitochondrial quality control and stress-induced remodeling, including *Rubcnl, Bmf*, and *Pmaip1*, were also elevated in the absence of AR in beige cells (Fig. 3A). Together, these transcriptomic alterations indicate that AR deficiency disrupts the metabolic program required for beige adipocyte mitochondrial function, while simultaneously inducing stress-response pathways.

Consistent with the transcriptional alterations identified by RNA-seq, we next examined whether AR deficiency affected mitochondrial morphology and organization in beige adipocytes. Transmission electron microscopy analysis of ingWAT from 10-week-old mice revealed profound mitochondrial abnormalities in AR^iB-/y^ adipocytes, characterized by enlarged mitochondria displaying matrix swelling, altered electron density, and disrupted cristae architecture compared with control adipocytes (Fig. 3B). Similar, although less pronounced, mitochondrial alterations were observed in BAT from AR^iB-/y^ mice (Fig. S3A).

To further characterize mitochondrial alterations following AR loss, we performed multiplex imaging analysis of mitochondrial markers in ingWAT, using Hyperion imaging mass cytometry. AR-deficient adipocytes displayed altered distribution of the mitochondrial outer membrane protein VDAC1, which exhibited a punctate pattern rather than the more continuous mitochondrial network observed in control adipocytes, together with increased expression of the mitochondrial fusion regulator OPA1 (Fig. 3C-3E). In parallel, increased expression of the antioxidant enzyme glutathione synthase (GSS) indicated activation of mitochondrial redox adaptation mechanisms in response to mitochondrial stress.

Given the accumulation of structurally altered mitochondria, we next investigated whether mitochondrial quality control pathways were engaged. Immunoblot analysis of ingWAT lysates revealed increased levels of mitophagy markers PINK1, PARKIN, and p62 in AR^iB-/y^ mice compared with controls (Fig. 3F). Together with the ultrastructural abnormalities, altered localization, and transcriptional induction of stress-response pathways, these findings indicate activation of mitochondrial quality control mechanisms in response to AR deficiency. However, despite observed mitochondrial defects, the body temperature, mitochondrial respiration, and relative hydrogen peroxide production were comparable between control and AR^iB-/y^ ingWAT samples at 10 weeks of age (Fig. S3B-E).

To determine whether AR loss impacts lipid accumulation and the mitochondria in a cell-autonomous manner in beige adipocytes, we differentiated immortalized cells derived from the stromal-vascular fraction from inguinal depot into mature beige adipocytes as described [20], and performed siRNA-mediated AR silencing. AR knockdown resulted in the accumulation of smaller lipid droplets and a fragmented mitochondrial pattern compared with the interconnected mitochondrial network observed in control cells (Fig. 3G and 3H). Consistent with the *in vivo* findings, RT-qPCR analysis in differentiated adipocytes demonstrated reduced expression of genes associated with mitochondrial metabolism (*Bnip3, Paox, Mfn1*) and adipocyte identity (*Pgc1a, Rxrg, Prdm16, Sorbs1*) following AR silencing (Fig. S3B). Together, these findings demonstrate that AR signaling is required to maintain mitochondrial metabolic competence and structural integrity in beige adipocytes.

### AR-deficient beige adipocytes remodel the immune microenvironment

Following the identification of mitochondrial stress and metabolic reprogramming in AR-deficient beige adipocytes, we next investigated whether impaired adipocyte homeostasis influenced the surrounding tissue microenvironment. Beyond metabolic pathways, differential expression analysis of ingWAT from AR^iB-/y^ mice revealed a strong enrichment of immune-associated pathways, including complement activation, innate immune responses, and phagocytic processes (Fig. 4A and S2G). Among the genes upregulated following AR deletion, several transcripts were associated with macrophage identity and activation (*Cd163, Cd52, Cd68, Cd74, Cd83, Cd86, Lyz2, Rac2*), complement cascade (*C1qa, C1qb, C1qc, C1ra, C1s1, C3, C6, C7*), and immune-related phagocytosis (*Fcer1g, Fcgr1, Fcgr2b, Fcgr3, Lgals3, Trem2, Tyrobp*) (Fig. 4A). Given that bulk RNA-seq reflects the transcriptional activity of multiple cellular populations, we next sought to determine the cellular origin of these immune signatures.

**Figure 4:**
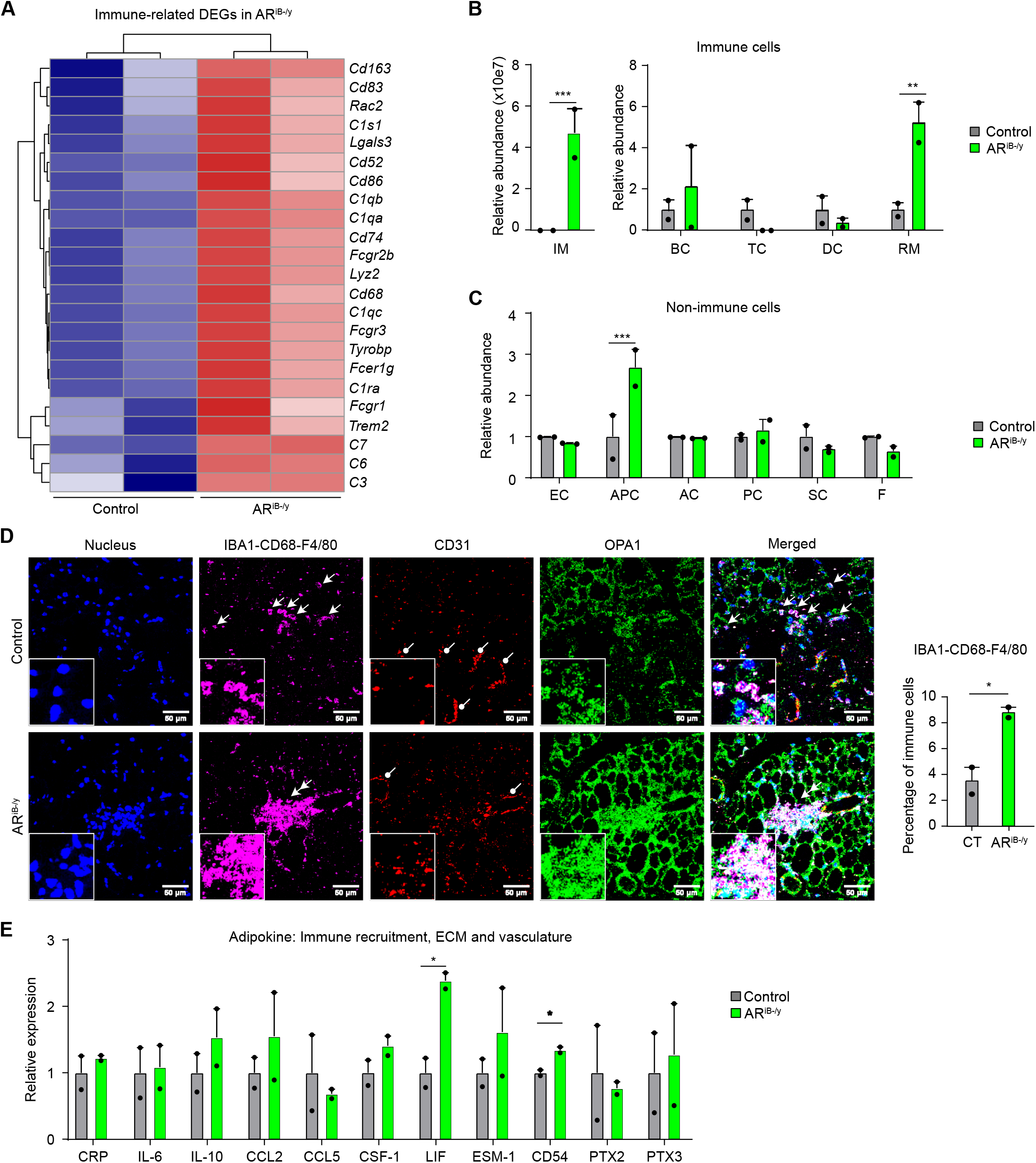
AR-deficient beige adipocytes remodel the immune microenvironment. (A) Heatmap depicting the mean-centered normalized expression of immune-associated genes obtained from RNA-eq datasets of ingWAT from control and AR^iB-/y^ mice at 10 weeks of age. (B-C) Relative abundance of the number of immune (B) and non-immune (C) cell populations in ingWAT of 10-week-old control and AR^iB-/y^ mice. IM = infiltrating macrophages; BC = B Cells; TC = T Cells; DC = Dendritic Cells; RM = Resident Macrophages; EC = Endothelial Cells; APC = Adipocyte Progenitors; AC = Adipocytes; PC = Pericytes; SC = Schwann cells; F = Fibroblasts. Data are presented as mean + SEM. Statistical test used is multiple two-tailed unpaired t-tests. **, p < 0.01; ***, p<0.001. (D)Imaging mass cytometry of IBA1-CD68-F4/80 (co-staining; magenta), CD31 (red), and OPA1 (green), and corresponding quantification of mean fluorescence intensity of ingWAT from 10-week-old control and AR^iB-/y^ mice. Nuclei are visible in blue. Insets at the lower-left corner represent higher magnification of corresponding images. Scale bars, 50⍰µm. Data are presented as mean + SEM. Statistical test used is two-tailed unpaired t-test. *, p < 0.05. (E)Quantification of selected adipokines relative expression secreted by ingWAT from 10-week-old control and AR^iB-/y^ mice. ECM = Extra-cellular matrix. Data are presented as mean + SEM. Statistical test used is two-tailed unpaired t-test. *, p < 0.05.

To this end, we performed deconvolution analysis of AR^iB-/y^ and control ingWAT transcriptomes using a published single-nucleus RNA-seq reference dataset from mouse ingWAT [21]. This analysis revealed a marked increase in macrophage-associated populations, including both resident and recruited macrophage subsets, as the major contributors to the immune-related transcriptional changes observed following AR deletion (Fig. 4B). In contrast, T cell populations were reduced, whereas adipocyte precursor populations were increased in AR^iB-/y^ ingWAT relative to control tissues, indicating selective remodeling of the immune-stromal compartment upon loss of AR signaling in beige adipocytes (Fig. 4C).

To validate macrophage accumulation at the tissue level and determine their spatial distribution, we performed imaging mass cytometry of key macrophage markers. Quantification of IBA1-, CD68-, and F4/80-positive cells demonstrated a significant increase in macrophage abundance in AR^iB-/y^ ingWAT compared with control tissues (Fig. 4D and S4A). Notably, macrophages were preferentially enriched in proximity to CD31-positive vascular structures (Fig. 4D), consistent with increased immune cell recruitment and vascular-associated tissue remodeling.

We next investigated whether paracrine signaling was altered in AR-deficient adipocytes, thereby contributing to macrophage recruitment. Analysis of adipose-derived inflammatory mediators revealed increased expression of CD54 (ICAM1) and LIF in AR^iB-/y^ ingWAT (Fig. 4E), two factors previously implicated in immune cell adhesion, activation, and macrophage recruitment. Together, these findings demonstrate that loss of AR signaling in beige adipocytes induces secondary remodeling of the immune microenvironment, characterized by macrophage accumulation and activation of complement and phagocytic programs. These immune alterations likely represent a consequence of impaired adipocyte homeostasis and mitochondrial/metabolic stress rather than a primary defect in immune regulation.

### AR deficiency induces long-term transcriptional adaptation but limits adipocyte metabolic flexibility

Having established that AR signaling is required for beige-to-white adipocyte transition during the early post-pubertal remodeling phase (Fig. 1A), we next investigated the impact of AR deficiency on long-term adipose tissue maturation. To determine whether the transcriptional repertoire impaired upon AR loss were maintained after endogenous whitening with age, we performed bulk RNA-sequencing analyses of ingWAT from 15-week-old control and AR^iB-/y^ mice. Differential expression analysis identified 699 differentially expressed genes (DEGs) in AR^iB-/y^ mice relative to control littermates (Fig. 5A). Compared with the extensive transcriptional response observed at 10 weeks of age (Fig. 2F), the number of differentially expressed genes was markedly reduced, indicating that AR-deficient adipocytes undergo progressive transcriptional remodeling during long-term tissue adaptation. In addition, integration of RNA-seq datasets obtained at 10 and 15 weeks of age identified a very limited subset of genes that remained persistently deregulated across time (Fig. 5B). Among the DEGs, downregulated genes were enriched for pathways associated with fatty acid biosynthesis and adipocyte differentiation (Fig. 5C). Representative genes persistently downregulated at both ages included Cebpd, *Irf4* and *Klf15* (Fig. 5D). Conversely, genes induced following AR loss in older mice were associated with signaling pathways of Insulin, FoxO, and calcium (Fig. 5E). Representative genes persistently upregulated at both ages included *Adcy7, Ccr2*, and *Ccr5* (Fig. 5D). Together, these findings indicate that, while the acute transcriptional response following AR deletion is progressively attenuated, AR-deficient adipocytes retain alterations in metabolic and stress-associated gene programs.

**Figure 5:**
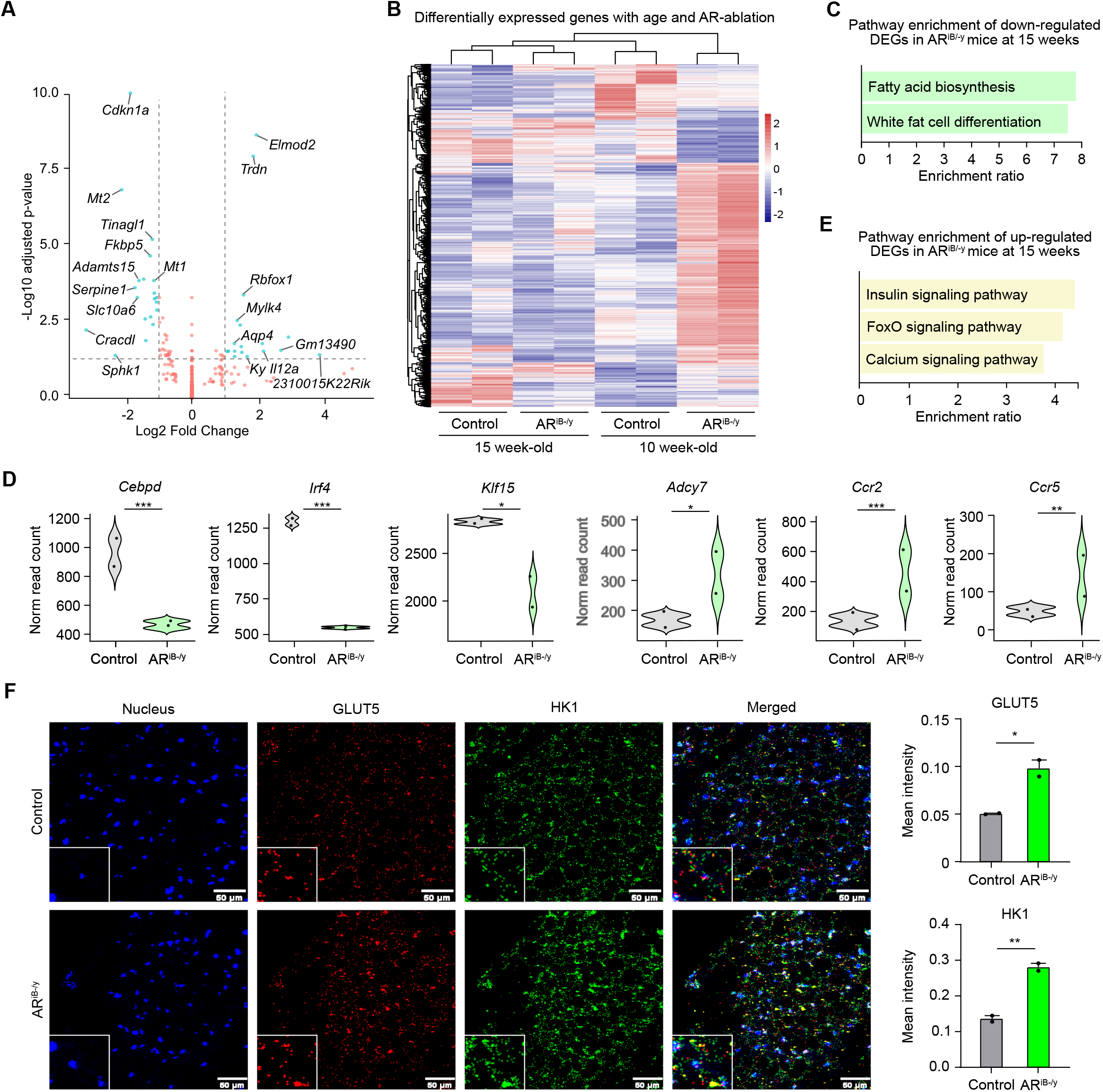
AR deficiency induces long-term transcriptional adaptation but limits adipocyte metabolic flexibility. (A)Volcano plot depicting in cyan the genes differentially expressed between AR^iB-/y^ and control ingWAT at 15 weeks. Selected markers are annotated. Additional genes are indicated in red. (B)Heatmap depicting the mean-centered normalized expression of differentially expressed genes obtained from the RNA-seq datasets of ingWAT from control and AR^iB-/y^ mice at 10 and 15 weeks of age. (C-E) Pathway enrichment of down-(C) and upregulated genes (E), and corresponding representative violin plots (D) for their normalized read counts, as obtained from RNA-seq data of ingWAT from 15-week-old control and AR^iB-/y^ mice. P-values were obtained from DESeq2. *, p < 0.05; **, p < 0.01; ***, p < 0.001. (F) Imaging mass cytometry of GLUT5 (red) and HK1 (green), and corresponding quantification of mean fluorescence intensity of ingWAT from 10-week-old control and AR^iB-/y^ mice. Nuclei are visible in blue. Inset at the lower-left corner of each image represent higher magnification of corresponding images. Scale bars, 501µm. Data are presented as mean + SEM. Statistical test used is two-tailed unpaired t-test.*, p < 0.05; **, p < 0.01

Considering that the differential transcriptome at 10 weeks and 15 weeks both include genes related to the glycolysis pathway, we assessed metabolic markers in ingWAT using imaging mass cytometry. Interestingly, while glycolysis pathway was attenuated at 10 weeks, AR-deficient ingWAT displayed increased expression of HK1 and the fructose transporter GLUT5 (Fig. 5F). These findings suggest that AR-deficient adipocytes undergo compensatory metabolic adaptation characterized by increased glycolytic activity, while failing to fully restore the transcriptional programs required for appropriate adipocyte maturation and lipid handling.

We next investigated whether this altered metabolic state affected the ability of AR-deficient adipocytes to adapt to an additional metabolic challenge. Control and AR^iB-/y^ mice were fed a high-fat diet (HFD) for 5 weeks following AR deletion. While HFD feeding induced the expected remodeling of control ingWAT, characterized by increased lipid droplet size and adipose tissue expansion, AR-deficient ingWAT displayed limited changes in lipid droplet distribution and impaired adipose tissue expansion (Fig. S5A-D). Together, these findings demonstrate that AR signaling is required not only for developmental beige-to-white adipocyte remodeling, but also for maintaining adipocyte metabolic flexibility during long-term tissue adaptation and nutrient excess.

### AR signaling is required for adaptive beige adipocyte remodeling during cold exposure

Following the identification of AR as a key regulator of beige adipocyte identity and metabolic function, we next investigated whether AR signaling was required for adaptive adipose tissue remodeling in response to physiological stress. To this end, control and AR^iB-/y^ mice were exposed to cold temperature (10°C) for 10 days, one week after AR deletion. Histological analysis of ingWAT revealed that cold exposure induced the expected remodeling of control adipocytes, characterized by increased abundance of small multilocular lipid droplets consistent with beige adipocyte activation (Fig. 6A). In contrast, AR^iB-/y^ ingWAT displayed limited morphological adaptation following cold exposure, with lipid droplet distribution remaining similar to that observed under room temperature conditions (Fig. 6A and Fig. S6A). These observations indicate that AR-deficient adipocytes exhibit impaired remodeling capacity in response to increased thermogenic demand.

**Figure 6:**
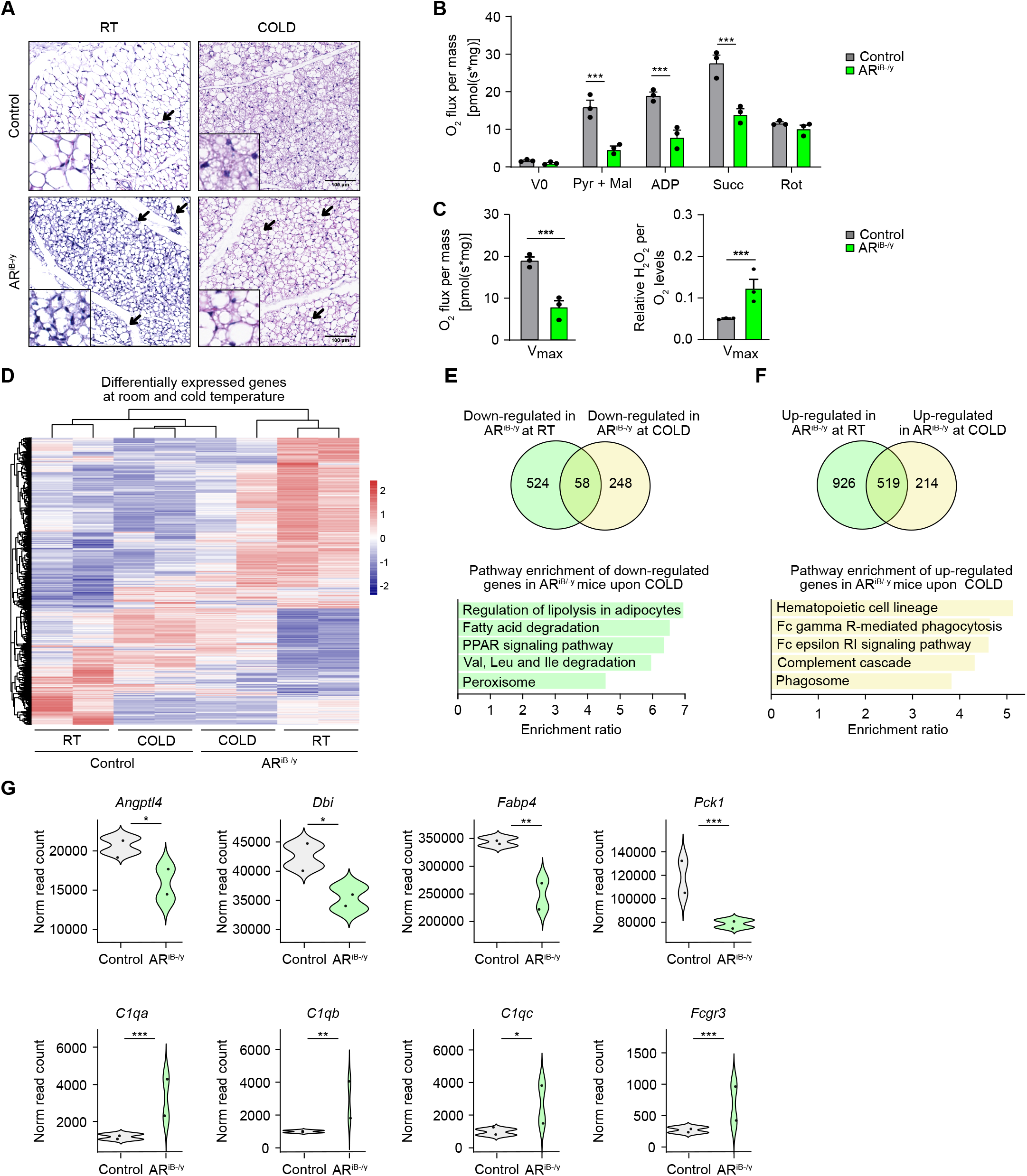
AR signaling is required for adaptive beige adipocyte remodeling during cold exposure. (A) Hematoxylin and Eosin (H&E) staining of ingWAT from control and AR^iB-/y^ mice maintained at room temperature (RT) or at 10°C for 10 days (COLD). Black arrows indicate dense lipid droplet clusters, representative of beige adipocytes. Insets at the lower-left corner represent higher magnification of corresponding images. Scale bars, 1001µm. (B-C) Quantification of mitochondrial oxygen (O_2_) flux per mass (B), and corresponding maximal respiration and relative maximal hydrogen peroxide (H2O2) flux per unit O2 flux (C) under successive supplementation of substrates for different mitochondrial complexes followed by inhibition of complex-I via Rotenone (Rot), in ingWAT of control and AR^iB-/y^ mice maintained at 10°C for 10 days. Data are presented as mean + SEM. Statistical test used is multiple two-tailed unpaired t-tests. ***, p<0.001. Pyr + Mal = Pyruvate + Malate; ADP = Adenosine Diphosphate; Succ = Succinate. (D) Heatmap depicting the mean-centered normalized expression of differentially expressed genes obtained from the RNA-seq datasets of ingWAT from control and AR^iB-/y^ mice maintained at room temperature (RT) or at 10°C for 10 days (COLD). (E) Overlap of the genes downregulated between AR^iB-/y^ and control mice at room temperature or with cold, and pathway analysis for genes which are downregulated in AR^iB-/y^ mice under cold exposure. (F) Overlap of the genes upregulated between AR^iB-/y^ and control mice at room temperature or with cold, and pathway analysis for genes which are upregulated in AR^iB-/y^ mice under cold exposure. (G) Violin plots for normalized read counts of representative genes differentially expressed in AR^iB-/y^ mice under cold exposure. P-values were obtained from DESeq2. *, p < 0.05; **, p < 0.01; ***, p<0.001.

To determine whether defective morphological adaptation was associated with altered mitochondrial function, we assessed mitochondrial respiration and oxidative stress markers following cold exposure. While mitochondrial respiration was comparable between genotypes under basal conditions (Fig. S3C-E), cold-exposed AR^iB-/y^ ingWAT displayed reduced maximal respiratory capacity and increased hydrogen peroxide production relative to cold-exposed controls (Fig. 6B and C). Together, these findings demonstrate that AR signaling is required for maintaining mitochondrial adaptability during cold-induced beige adipocyte remodeling.

To further define the transcriptional mechanisms underlying impaired adaptation, we integrated transcriptomic datasets obtained from AR-deficient ingWAT under basal conditions at 10 weeks and after cold exposure. Differentially expressed genes identified at 10 weeks (AR^iB-/y^ versus control ingWAT; 1,477 upregulated and 673 downregulated genes) were used a reference signature of the early transcriptional consequences of AR loss. Bulk RNA-seq analyses of ingWAT from cold-exposed mice identified 939 genes significantly altered in AR^iB-/y^ mice relative to cold-exposed control littermates (Fig. S6B). Projection of the AR-dependent signature under endogenous conditions onto cold-exposed mice revealed distinct temporal trajectories of gene regulation (Fig. 6D). Among the DEGs, downregulated genes were enriched for pathways associated with adipocyte metabolism, including lipid metabolism, PPAR signaling, and peroxisomal function (Fig. 6E). Representative genes persistently downregulated under both room temperature and cold included Angptl4, Dbi, Fabp4, and Pck1 (Fig. 6G), which are associated with PPAR signaling pathway. Conversely, genes induced following AR loss and maintained or further enhanced during cold exposure were associated with stress responses and immune-related pathways, including complement activation and phagocytosis programs (Fig. 6F). Representative genes persistently upregulated under both room temperature and cold included *Fcgr1, Fcgr3, C1qa, C1qb*, and *C1qc* (Fig. 6G). These findings indicate that although AR-deficient adipocytes undergo partial transcriptional compensation over time, they fail to restore the metabolic flexibility required during increased physiological demand. Together, these results demonstrate that AR signaling establishes a transcriptional framework required for adaptive beige adipocyte remodeling. Loss of AR leads to a progressive impairment of metabolic plasticity, characterized by defective mitochondrial adaptation and persistent activation of secondary stress and immune-associated programs.

## DISCUSSION

Adipose tissue remodeling requires coordinated changes in adipocyte identity, metabolism, and interactions with the surrounding tissue microenvironment. Although androgen receptor (AR) signaling has long been implicated in adipose tissue biology and adipogenesis, its specific contribution to the maintenance and remodeling of mature beige adipocytes has remained largely unexplored. Here, we identify AR as a central regulator of post-pubertal beige adipocyte remodeling in male mice. By combining inducible genetic ablation, cistromic and transcriptomic analyses, and physiological challenges, we demonstrate that AR establishes a transcriptional program required to preserve metabolic competence and adaptive plasticity of mature beige adipocytes. Importantly, our data reveal that morphological persistence of beige adipocytes following AR deletion does not reflect preservation of their functional identity, thereby uncoupling beige morphology from beige adipocyte function.

This distinction has important implications for our understanding of beige adipocyte biology. Beige adipocytes are generally identified by their multilocular morphology and expression of thermogenic markers, yet these features are often assumed to directly reflect metabolic activity [22, 23]. In our model, AR-deficient adipocytes retained characteristic multilocular lipid droplets and failed to complete the physiological beige-to-white transition during aging. Surprisingly, this apparent maintenance of beige morphology was accompanied by impaired mitochondrial integrity, altered metabolic gene expression, and defective adaptation to cold exposure. These observations indicate that AR is not required to generate or morphologically maintain beige adipocytes per se, but rather to preserve their metabolic competence. Our findings therefore indicate that beige adipocyte morphology and beige adipocyte function constitute two separable biological properties, highlighting the existence of an intermediate dysfunctional beige state that remains morphologically beige despite profound metabolic impairment. Previous studies have established multiple roles for AR signaling in adipose tissue biology, including the regulation of adipocyte differentiation, lipid metabolism, and systemic energy homeostasis [24]. More recently, AR was proposed to function as a negative regulator of beige adipocyte formation through repression of PRDM16-dependent transcriptional programs [25]. Using constitutive adipocyte-specific AR deletion and androgen stimulation approaches, this study showed that reduced AR activity promotes beige adipocyte features, including increased expression of thermogenic genes and enhanced mitochondrial respiration. Several observations are consistent between these findings and our study, including the depot-specific expression pattern of AR and the lack of detectable AR occupancy at the Ucp1 locus. However, the functional consequences of AR deletion differ substantially between the two experimental models. While constitutive AR deletion increased thermogenic gene expression, oxidative capacity, and beige adipocyte characteristics, our inducible *Ucp1*-CreER^T2^-mediated ablation of AR in established beige adipocytes resulted in reduced thermogenic gene expression, impaired mitochondrial function, increased oxidative stress, and defective adaptation to cold exposure. These divergent outcomes may reflect differences in the biological context in which AR function was interrogated. The Adipoq-Cre model used previously induces AR deletion across the adipocyte compartment from early stages of adipocyte development, thereby integrating potential effects on adipocyte differentiation, recruitment, and subsequent remodeling. In contrast, our inducible *Ucp1*-CreER^T2^ strategy selectively targets pre-existing beige adipocytes at adult stage after their establishment, allowing us to specifically examine AR-dependent mechanisms required for maintenance of mature beige adipocyte function. Additional differences, including pharmacological AR activation by dihydrotestosterone versus analysis of endogenous AR activity, exon targeting strategies, and *ex vivo* versus *in vivo* genomic approaches, may further contribute to the distinct transcriptional and physiological outcomes observed. Together, these studies indicate that AR regulates multiple layers of beige adipocyte biology, ranging from control of beige adipocyte programming to preservation of metabolic fitness and adaptive capacity in mature beige adipocytes.

Our genomic analyses further support this model. Integration of AR ChIP-seq with RNA sequencing identified AR as a regulator of enhancer-associated transcriptional programs governing adipocyte identity, lipid metabolism, peroxisomal function, and mitochondrial homeostasis. Notably, we did not detect AR occupancy at the Prdm16 or Ucp1 loci, suggesting that AR does not directly regulate canonical beige lineage determinants in mature adipocytes. Instead, AR preferentially binds distal regulatory elements associated with genes such as *Fabp4, Lpl, Adipoq, Adrb3, Aqp7*, and *Slc7a10*, supporting a model in which AR maintains the metabolic architecture of mature beige adipocytes rather than specifying their lineage. This interpretation is further supported by the predominance of canonical androgen response elements within AR-binding sites and by the strong enrichment of metabolic pathways among putative direct AR targets.

The metabolic defects observed following AR ablation were accompanied by profound alterations in mitochondrial organization and quality control. Mitochondria from AR-deficient beige adipocytes displayed swelling, disrupted cristae organization, accumulation of PINK1/PARKIN proteins, and elevated oxidative stress following cold exposure. Rather than representing a primary consequence of impaired mitochondrial clearance known to be associated with beige-to-white transition, these alterations are likely secondary to the extensive metabolic rewiring induced by AR loss. Indeed, mitochondrial quality control pathways are highly sensitive to changes in cellular metabolism, and stabilization of PINK1 protein occurs primarily through impaired mitochondrial import following membrane depolarization rather than increased transcription [26-28]. This is consistent with our observation that mitophagy-associated proteins accumulated despite relatively modest transcriptional alterations. Together, these findings place mitochondrial dysfunction downstream of AR-dependent transcriptional remodeling and suggest that defective mitochondrial quality control contributes to the loss of beige adipocyte fitness.

An additional consequence of AR deficiency was extensive remodeling of the adipose tissue immune microenvironment. Transcriptomic deconvolution, imaging mass cytometry, and cytokine profiling consistently indicated increased macrophage recruitment following AR deletion. Rather than reflecting a direct effect of AR loss on immune cells, which do not undergo recombination in our model, these changes most likely arise secondarily to adipocyte dysfunction. Metabolically stressed adipocytes are known to secrete chemotactic molecules capable of recruiting circulating monocytes and macrophages [29, 30], and our identification of increased ICAM1 and LIF expression supports this mechanism. Interestingly, macrophages were frequently observed in close proximity to the adipose vasculature before becoming more diffusely distributed throughout the tissue, suggesting active recruitment from the circulation rather than local expansion. Whether these recruited macrophages ultimately contribute to tissue repair, lipid clearance, or further metabolic dysfunction remains an important question for future studies.

Finally, our analyses under aging and cold exposure emphasize that the physiological role of AR extends beyond maintenance of basal adipocyte homeostasis to the preservation of adaptive plasticity. Although transcriptional compensation partially emerged during aging, AR-deficient adipocytes failed to appropriately respond to cold-induced thermogenic demand. Cold adaptation involves both reactivation of pre-existing dormant beige adipocytes and de novo differentiation of beige adipocytes from precursor populations, with their relative contribution depending on prior cold exposure history [31]. Because Ar deletion in our model is restricted to UCP1-expressing adipocytes, our findings primarily demonstrate that AR is required for the adaptive remodeling of pre-existing beige adipocytes rather than their de novo formation. The inability of AR-deficient adipocytes to restore metabolic function under cold challenge therefore reveals that morphological persistence alone is insufficient to support physiological thermogenic adaptation.

Collectively, our findings redefine the role of androgen receptor signaling in beige adipocyte biology. Rather than functioning as a simple positive or negative regulator of beige adipocyte identity, AR establishes a transcriptional framework that preserves metabolic competence, mitochondrial fitness, and adaptive remodeling. Loss of AR uncouples beige adipocyte morphology from function, trapping adipocytes in a metabolically compromised intermediate state that ultimately reshapes the surrounding tissue microenvironment. More broadly, our work highlights the importance of distinguishing cellular identity from cellular function when investigating adipose tissue plasticity and identifies AR signaling as a key determinant of beige adipocyte fitness during post-pubertal remodeling.

## METHODS

### Mouse studies

All experiments were performed in an accredited animal house, in compliance with French and EU regulations on the use of laboratory animals for research. Intended manipulations were approved by the Ethical committee (Com’Eth, Strasbourg, France) and authorized by the French Research Ministry (MESR), conforming to the 2010/63/EU directive (decision number APAFIS#23909-2020010713409223 v8). Mice were maintained in a temperature (19-23°C) and humidity-controlled (40-60%) animal facility with a 12-h light/dark cycle. Standard rodent chow (2800⍰kcal/kg, Usine d’Alimentation Rationelle, Villemoisson-sur-Orge, France) and water were provided ad libitum.

For high-fat diet experiments, mice were maintained at aforementioned temperature with a 12h light/dark cycle, free access to water, and modified rodent diet (60 kcal % fat and 1.5x vitamin mix, Research Diets Inc., New Brunswick, NJ, USA) for 5 weeks. For cold exposure experiments, mice were maintained at 10 °C with a 12h light/dark cycle, free access to water, and standard rodent chow for 10 consecutive days. Body temperature was measured with a rectal probe linked to a digital thermometer (Bioseb).

Mice were euthanized by cervical dislocation, and tissues were immediately harvested, weighed, and either frozen in liquid nitrogen or processed for biochemical and histological analysis. All harvested ingWAT tissues used for non-histological experiments had their residing lymph node removed and only part of the remaining tissue circling around the lymph node was utilized for further analysis. Nomenclature for dissected adipose depots was used as described previously [32]

### Generation of Conditional AR-KO Mice

For beige adipocyte-specific androgen receptor (AR) ablation, AR^L2/L2^ female mice, carrying LoxP sites flanking chromosome X exon 1 encoding the AR N-terminal domain [33], were intercrossed with *Ucp1*-CreER^T2^ male mice [5] to generate *Ucp1*-CreER^T2^/AR^L2/Y^ mutant mice, which express CreER^T2^ recombinase selectively in brown and beige adipocytes. To induce recombination, 9-week-old AR^L2/Y^ control males and their *Ucp1*-CreER^T2^/AR^L2/Y^ pre-mutant littermates received intraperitoneal tamoxifen injections (1 mg/mouse/day) for five consecutive days, generating control and AR^(i)B−/Y^ mutant mice. All mice were on a C57Bl/6J background. Primers used for genotyping are listed in Supplementary Table 1.

### RNA extraction and analysis

Adipose tissue was homogenized in TRIzol reagent (TRIREAGENT, Molecular Research Center, TR-118, 513-841-0900). RNA was isolated using a standard phenol/chloroform extraction protocol, and quantified by spectrophotometry (Nanodrop, Thermo Fisher Scientific). 1⍰µg of total RNA underwent reverse transcription using SuperScript IV (Life Technologies) with oligo(dT) primers, according to the supplier’s protocol. cDNA was diluted 100 times and quantitative PCR (qPCR) was performed with a Lightcycler 480 II (Roche) using the SYBR® Green PCR kit (Roche) according to the supplier’s protocol (2⍰µl cDNA, 4.8⍰µl H_2_O, 5⍰µl Syber Green 2x mix and 0.2⍰µl of 100⍰µM primer mix). Primers are described in Supplementary Table 2. 18S primers were used as internal control. Data were analyzed using the ΔΔCt method [34].

For RNA-seq, RNA integrity was confirmed by Bioanalyzer. The cDNA library was prepared at the GenomEast platform from IGBMC, and sequenced with the standard Illumina protocol (NextSeq 2000-X, single-end, 50⍰bp) following the manufacturer’s instructions. Removal of adapter, polyA and low-quality sequences (Phread quality score below 20) were done using cutadapt 4.2. FastQC 0.12.1 [35] was used to evaluate the quality of sequencing. Reads were mapped to the GRCm39 assembly of *Mus musculus* genome (Gencode version M38) using STAR 2.7.11a [36], and the BAM files generated were indexed using samtools 1.21 [37]. Only uniquely aligned reads were retained for further analyses. Gene raw count matrix was obtained using Subread 2.0.6 [38]. For comparison among datasets, transcripts with more than 10 raw reads were considered. Differentially expressed genes (DEGs) were identified using the Bioconductor library DESeq2 [39] with a *p-value*⍰<⍰0.05. Pathway analysis was done using WebGestalt using the Over-Representation Analysis (ORA) method and a false discovery rate (FDR)1<10.05 as the significance threshold [40]. Heatmaps of normalized expression values were generated with R. Genes were clustered according to the hierarchical method (HCL clustering) using pheatmap in R [41]. Transcription factor predictions were done using LISA [42].Overlap of our RNA-Seq data with single-nuclei RNA-Seq from literature was done using the DWLS algorithm [43]. Venn diagrams were generated using Venny [44].

### Chromatin Immunoprecipitation Sequencing (ChIP-Seq)

Chromatin Immunoprecipitation was performed on whole inguinal white adipose tissue extracts as described [45], using 10⍰µg anti-AR (AB 108341; Abcam) and anti-H3K4me2 (39141; Active Motif), along with rabbit IgG (Santa Cruz, sc2357) negative control bound to protein Dynabeads protein G (Thermo Fisher Scientific, 10004D). DNA libraries were prepared and sequenced with an Illumina NextSeq 2000-X (paired-end, 50⍰bp reads) as per the manufacturer’s instructions. FastQC 0.11.5 [35] was used to evaluate the quality of sequencing. Reads were and mapped to the mm10 reference genome using Bowtie 1.3.1 [46]. Uniquely mapped reads were retained for further analysis. Reads overlapping with ENCODE hg38 blacklisted region V2 were removed using Bedtools [47]. Bigwig files were generated using Homer software makeUCSCfile script with default parameters and scaled to 1e7 reads [48]. MACS2 (2.2.9.1) algorithm was used for the peak calling [49]. All peaks with an FDR greater than 0.05 were excluded from further analysis. The genome-wide intensity profiles were visualized using the IGV genome browser [50]. HOMER was used to annotate peaks and for motif searches [48]. Genomic features (promoter/TSS, 5’ UTR, exon, intron, 3’ UTR, TTS and intergenic regions) were defined and calculated using Refseq and HOMER according to the distance to the nearest TSS. Clustering analyses were done with the deeptools software [51], and clustering normalization was done with the K-Means linear option. Pathway analysis was performed with WebGestalt using the Over-Representation Analysis (ORA) method [40]. Overlap of our RNA-Seq data with single-nuclei RNA-Seq from literature was done using the DWLS algorithm. Gene centric analysis was performed with windowbed from bedtools.

Parameters were set as default, with the exception of the following: Bowtie (-m 1 -- strata -- best - y - S - l 40), MACS2 [callpeak -- gsize 1.87e9 -- create_model -- extsize 300 -- nobroad -- keep-dup all], deeptools (input bed files normalized to 20 million reads per sample).

### snRNAseq analyses

PRJNA1010853 was used for snRNA-seq analysis of mouse ingWAT at 10 weeks of age, using metacluster annotations, as described [21]. Bioinformatics analyses were assessed with Seurat v5.0.1. DWLS (dampened weighted least squares) algorithm was used to deconvolute bulk RNA-Seq dataset using the snRNA-seq dataset. Data normalization was performed with -NormalizeData, and scaled with -ScaleData with default features.

### Mitochondrial Respiration

Inguinal adipose tissues were isolated, immersed immediately in KHB buffer (120 mM NaCl, 4.7 mM KCl, 1.18 mM KH2PO4, 1.17 mM MgSO4, 1.8 mM CaCl2, 30 mM Na-HEPES, 1 mM NaHCO3 (added fresh), 1% (w/v) fatty acid–free BSA (Sigma), pH adjusted to 7.4 with NaOH), and homogenized on ice. The homogenate was moved to the chambers of O2k (OROBOROS instruments, Innsbruck, Austria) to detect O2 flux from mitochondria in presence of different substrates namely pyruvate, malate, ADP, succinate, and rotenone as described [52]. The O2 and H2O2 flux were monitored on the Oroboros Datlab software (OROBOROS instruments, Innsbruck, Austria).

### Histological and Immunofluorescence Analysis

Adipose tissue was fixed in 4% PFA overnight at 4 °C then washed in PBS and 70% ethanol and embedded in paraffin. 5 μm thick paraffin sections were cut at room temperature (RT) using a microtome (Leica RM, 2145, 0737/09.1998) and stored at 4 °C until further analysis.

For hematoxylin and eosin (H&E) staining, paraffin sections underwent deparaffinization and rehydration, followed by staining with Papanicolaou’s solution 1a Harris’ Hematoxylin solution (Sigma-Aldrich ref 1092530500) for 3 min. Afterwards, the slides were washed and stained with 0.2% Eosin-Y for 3 min. The slides were then washed, dehydrated, and mounted with PERTEX mounting medium.

For both immunohistochemistry (IHC) and immunofluorescence (IF), paraffin sections underwent deparaffinization and rehydration, followed by antigen retrieval under controlled temperature and pressure conditions for 20⍰min in Signal Stain® Citrate Unmasking Solution (Cell Signalling 14746S). For IHC, endogenous peroxidase activity was quenched with peroxide incubation for 10 min, followed by blocking for 1 hour at room temperature. Afterwards, the sections for both IHC and IF were incubated overnight at 4⍰°C with the primary antibodies directed against AR (Rabbit, Abcam, AB108341, 1:200), UCP1 (Mouse, Invitrogen, GT3111, 1:200) and PLIN1 (Rabbit, Abcam, AB3526. 1:200), diluted in PBST. Mouse (Santa Cruz Biotechnology, sc-2025, 1:500) or rabbit (PeproTech, 500-P000-500UG, 1:500) normal IgGs were used as controls. Sections were rinsed with PBS, and incubated with appropriate Alexa-Fluor-conjugated secondary antibodies with different fluorescent wavelengths (488 or 555 nm, 1:500, Invitrogen: a32732, a32723) in PBST for 45 min at RT. For IF, the slides were washed and then mounted with Fluoromount™ Aqueous Mounting medium containing DAPI (Invitrogen, 00-4959-52). For IHC, the slides were washed and incubated with DAB detection stain (SignalStain DAB Substrate Kit #8059S) for required duration, followed by Hematoxylin staining.

Cross-Sectional Area (CSA) measurements for lipid droplets in PLIN1 immunofluorescence-stained tissue sections was performed using CellPose [53] tool on QuPath [54].

### Adipose ultrastructure analysis

One mm2 adipose tissue samples were fixed overnight at 4°C by immersion in a mix of 2.5% glutaraldehyde and 2.5% paraformaldehyde diluted in 0.1 M cacodylate buffer (pH 7.4), washed in the same buffer for 30 min, and stored at 4°C. Post-fixation was performed with 1% osmium tetroxide in 0.1 M cacodylate buffer for 1 h at 4°C, followed by dehydration through a graded ethanol series (50%, 70%, 90%, and 100%) and propylene oxide (30 min each). Samples were oriented longitudinally or transversally and embedded in Epon 812. 70 nm ultrathin sections were obtained and contrasted with uranyl acetate and lead citrate. Imaging was performed at 70 kV using a Morgagni 268D electron microscope, with digital acquisition via a Mega View III camera (Soft Imaging System).

### Imaging Mass Cytometry Antibodies and Metal Conjugation

Antibodies were conjugated to the Cadmium isotope using the Maxpar® MPC9 Antibody Labeling Kit. All other antibodies were labeled using the Maxpar® X8 Antibody Labeling Kit according to the manufacturer’s instructions (PRD002 Rev 14, Fluidigm, Standard Biotools). List of antibodies used for the experiments are mentioned in Supplementary Table 3.

### Tissue labeling

After deparaffinization and antigen retrieval using Dako Target Retrieval Solution at pH 9 (S236784-2, Agilent technologies) in a water bath (96°C for 30 min), 3 µm tissue sections were encircled with a Dako Pen, incubated with Superblock™ (37515, ThermofisherScientific) at room temperature (RT) for 45 min, and then with FcR Blocking Reagent (130-092-575, Miltenyi) at RT for 1h. After three washes (8 min/each) in PBS/0.2% Triton X-100 (PBS-T), metal-tagged antibodies (Supplementary Table 3) were diluted in PBS/1% BSA buffer. After incubation with the primary antibodies at 4°C overnight, sections were washed in PBS-T three times (8min/each) and nuclei were stained with iridium (1:400 in PBS; Fluidigm, Standard Biotools), a DNA Intercalator, for 30 min at RT. Sections were washed in PBS for 5 min, then in distilled water for 5 min, and dried at RT for 30 min.

### Data acquisition

Images were acquired with the Hyperion Imaging System (Fluidigm, Standard Biotools) according to the manufacturer’s instructions. After choosing the region of interest (ROI) in the section, the ROI was ablated with a UV laser at 200 Hz. Data were exported as MCD files and visualized using the Fluidigm MCD™ viewer 1.0.560.6. The minimum and maximum thresholds were adapted for each marker and for each tissue for optimal visualization. Gamma was set to 1.

### Protein Analyses

Adipose tissues were mechanically ground in RIPA buffer [50⍰mM Tris (pH 7.5), 1% NP-40, 0.5% sodium deoxycholate, 0.1% SDS, 150⍰mM NaCl, 5⍰mM EDTA pH 8.0, and a protease inhibitor cocktail (45⍰mg/ml; Roche, Cat. No. 11 873 580 001)] at 4⍰°C. Tissue lysates were centrifuged at 12,000⍰g for 10 min at 4⍰°C, and the supernatants were then collected for further analysis. For western blot analyses, homogenates were separated in 8% polyacrylamide gels, transferred to Hybond nitrocellulose membranes (Amersham Biosciences), and probed overnight at 4⍰°C with specific antibodies targeting AR (Abcam, AB108341, 1:500), UCP1 (Invitrogen, GT3111, 1:1000), PINK1 (Proteintech, 23274-1-AP, 1:1000), PARKIN (Proteintech, 14060-1-AP, 1:1000), P62 (Abcam, AB56416, 1:1000) and α-TUBULIN (IGBMC, 1Tub2A2, 1:1000). After washing, membranes were probed with secondary antibodies (anti-Mouse-HRP, Cell Signalling, 7076S, 1:5000; anit-Rabbit-HRP, Cell Signalling, 7074S, 1:5000) conjugated to horseradish peroxidase. For adipokine array, the adipose tissues were lysed and processed using the Proteome Profiler Mouse Adipokine Array Kit (R&D Systems #ARY013) as per manufacturer’s protocol. Membranes for both western blot and adipokine array were detected using chemiluminescence detection systems (GE Healthcare, Amersham, ImageQuant 800 or Imager 600). Protein quantification was assessed by the FIJI/ImageJ distribution software [55].

### Cell culture

Immortalized cell lines derived from the stromal-vascular fraction from inguinal depot of 129SVE mice were obtained from Dr. Anastasia Georgiadi’s laboratory. Cells were grown in Dulbecco’s Modified Eagle Medium/Nutrient Mixture F-12 (DMEM/F-12; ThermoFischer Scientific Cat# 10565018), supplemented with GlutaMAX, 10% fetal calf serum (FCS), along with 1% Penicillin-Streptomycin (PS). Cells were maintained at 37°C in a humidified atmosphere with 5% CO2. Cells were split at 70-80% confluence, and the medium was replaced every 48 hours.

Plates intended for cell differentiation were pre-coated with rat-tail collagen I. To induce differentiation into beige adipocytes, preadipocytes were seeded at 60% confluence and switched at confluence to differentiation medium 1 composed of DMEM/F-12 GlutaMAX, supplemented with 10 ^μ^M Rosiglitazone (SIGMA-ALDRICH, R2408), 5 ^μ^M Dexamethasone (SIGMA-ALDRICH, D1756), 0.5 mM isobutyl methylxanthine (IBMX, SIGMA-ALDRICH, I5879), 0.5 µg/mL Insulin (SIGMA-ALDRICH, I9278), 1 nM 3,3⍰,5-Triiodo-L-thyronine (T3, SIGMA-ALDRICH, T2752), for 3 days, followed by differentiation medium 2 made of DMEM/F-12 GlutaMAX, supplemented with 10 ^μ^M Rosiglitazone (SIGMA-ALDRICH, R2408), 0.5 µg/mL Insulin (SIGMA-ALDRICH, I9278), 1 nM 3,3⍰,5-Triiodo-L-thyronine (T3, SIGMA-ALDRICH, T2752) for 4 days, with medium changes every 48 hours.

For AR silencing experiments, differentiated beige adipocytes were transfected for 24 h with 30 nM of siRNA (Stealth, Thermo Scientific) against AR (forward: 5’-AGGAAUUCCUGUGCAUGAAAGCACU-3’; reverse: ^5’-AGUGCUUUCAUGCACAGGAAUUCCU-3’^) or scramble control siRNA (forward: 5’-CAAAGGAGUUCCACAGCUAUGGCUA-3’; reverse: 5’-UAGCCAUAGCUGUGGAACUCCUUUG-3’) diluted in Opti-MEM (Gibco, USA) using Lipofectamine™ RNAiMAX (ThermoFischer, 13778075) transfection reagent, following the manufacturer’s protocol. Cells were subsequently harvested for downstream processing.

For mitotracker assay, differentiation medium 2 was replaced with 200 nM Mitotracker Green FM (ThermoFisher Scientific M46750) diluted in PBS and cells were incubated for 30 min. Live imaging performed using Nikon Spinning Disc microscope.

## Data Analysis

Data analyses were conducted using DESeq2, Microsoft Excel and GraphPad Prism 9. In all graphical representations, error bars indicate the standard error mean (SEM). Comparisons between groups were performed using appropriate statistical tests, selected based on the number of groups, data distribution (normality), variance homogeneity, and multiple comparisons. The specific statistical tests applied are indicated in the figure legends. Statistical significance was defined as follows: ns (non-significant), p ≥ 0.05; *, p < 0.05; **, p < 0.01; and ***, p < 0.001

## Data Availability

All sequencing datasets generated in this study are deposited in the Gene Expression Omnibus (GEO) with the accession number GSE341702 for ChIP-seq analyses and GSE341704 for RNA-seq experiments.

## Acknowledgments

We thank Prof. C. Wolfrum (Helmholtz Diabetes Center) for generously providing the Ucp1-CreER^T2^ mouse line, and Dr. Anastasia Georgiadi for immortalized beige adipocytes. We are grateful to Nikola Djordjevic, Anastasia Bannwarth, Brayann Weis, and Régis Lutzing for their valuable technical support, and Tao Ye, Matthieu Jung, Céline Keime, and Pierrick Dupre for excellent bioinformatics assistance. We acknowledge the support of the IGBMC and Mouse Clinical Institute (ICS, Illkirch-Graffenstaden, France) core facilities, including Michaël Gendron, Elise Le Marchand, Alexandre Vincent, and Sylvie Falcone from the animal facility, Erwan Grandgirard and Simon Langlois from microscopy service, Nadia Messaddeq from the electron microscopy platform, the cell culture service, Olivia Wendling and Hugues Jacobs from histopathology services, as well as the flow cytometry service and GenomEast facility, a member of the ‘France Génomique’ consortium (ANR-10-INBS-0009). We are grateful to Daniela Rovito, Isabelle M. Billas, Elodie Montsellier, Karan Joshi and Mariia Nazarova for helpful discussions.

## Funding

This work was supported by the Interdisciplinary Thematic Institute IMCBio as part of the ITI 2021-2028 program of the University of Strasbourg, the Centre National pour la Recherche Scientifique (CNRS), Institut national de la santé et de la recherche médicale (Inserm), from IDex Unistra (ANR-10-IDEX-0002), SFRI-STRAT’US (ANR 20-SFRI-0012) and EUR IMCBio (ANR-17-EURE-0023) projects, under the framework of the French Investments for the Future Programme. Additional support was provided by Inserm, CNRS, Unistra, IGBMC and Agence Nationale de la Recherche (ANR-24-CE14-0247-01). R.S. was funded by IMCBio, S.S.C. by the ANR (MYOGLUCO, ANR-22-CE11-0014-01). J.G.R. received funding from the Programme CDFA-07-22 from the Université franco-allemande and Ministère de l’Enseignement Supérieur de la Recherche et de l’Innovation, E.C. by Ministère de l’Enseignement Supérieur de la Recherche et de l’Innovation.

## Author contributions

D.D. formulated the initial hypothesis. R.Sa., A.B., M.O., A.C., J.G.R., Q.C., S.S.C. and E.C. carried out functional, molecular, and histological experiments. D.D., R.Sa. and T.Y. executed bioinformatics analyses. R.Sa. and N.M. took responsibility of imaging analyses. R.Sa., P.D., L.L.-C. and M.L. took responsibility for Imaging Mass Cytometry analysis. E.M. and R.Sc. provided scientific material and assistance. R.Sa., D.M., and D.D. took primary responsibility for data analysis and writing the manuscript.

## Declaration of Interests

None declared.

## Declaration of generative AI and AI-assisted technologies

None declared.

## Notes

### Competing Interest Statement

The authors have declared no competing interest.

### Summary of Updates

Update in the author list in the main manuscript.

