## Supplementary Figures and Tables for "Androgen Receptor mediates beige adipocyte homeostasis and plasticity"

This file includes Supplementary Figures 1-6 and Supplementary Tables 1-3.

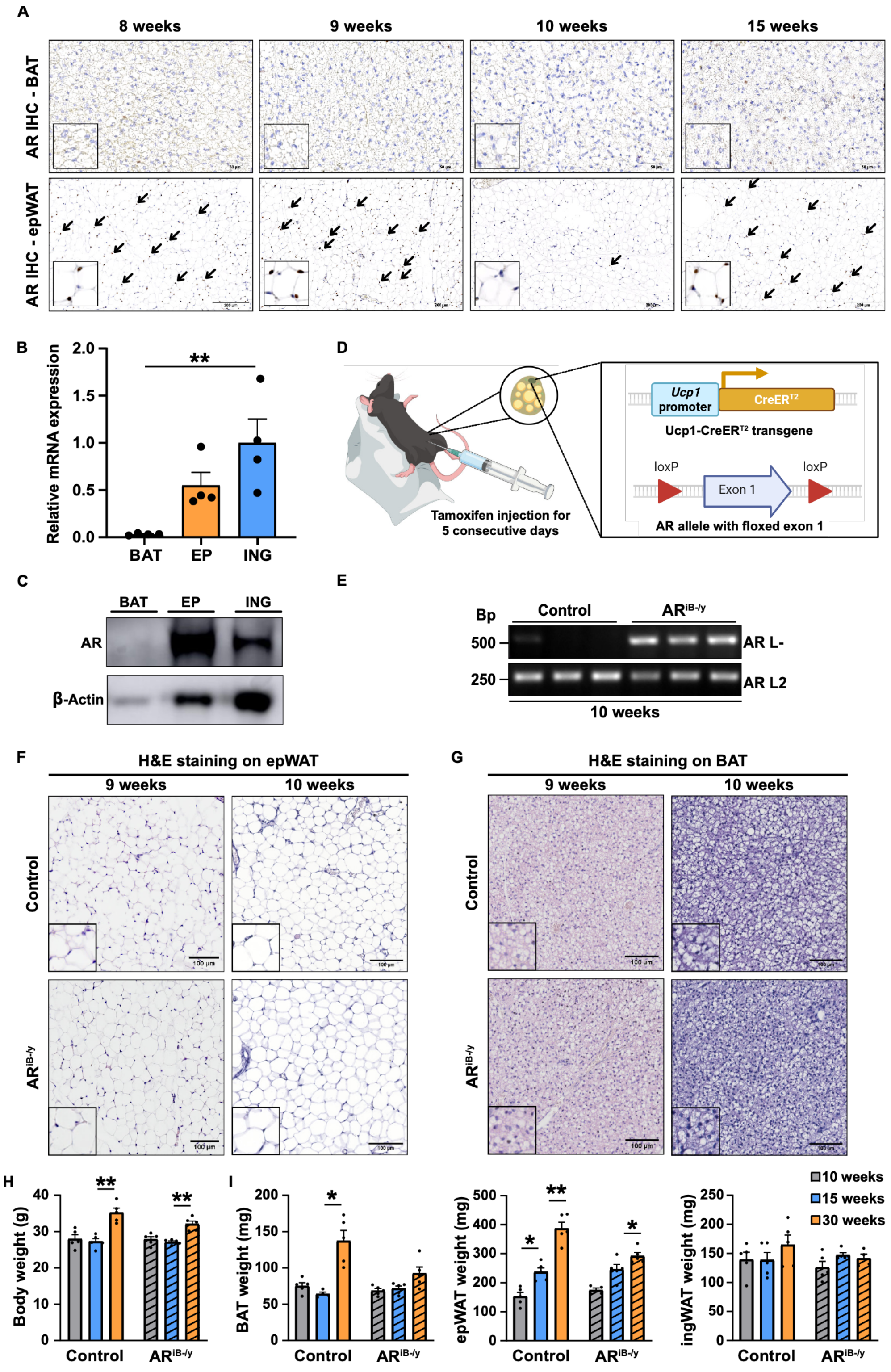


**Figure S1: AR expression in mouse adipose tissue and physiological consequences of its invalidation in beige adipocytes**

**(A)** Immunohistochemical (IHC) detection of the androgen receptor (AR) in brown adipose tissue (BAT) and epidydimal white adipose tissue (epWAT) of control mice at 8, 9, 10, and 15 weeks of age. Black arrows indicate AR expression in the nuclei of epWAT. Insets at the lower-left corner represent a higher magnification of corresponding images. Scale bars, 50 µm for BAT and 200 µm for epWAT.

**(B)** AR relative transcript levels in various fat pads of 10-week-old control mice. Data are presented as mean + SEM. Statistical test used was an ordinary one-way ANOVA. **, $p < 0.01$. ING = Inguinal WAT; EP = Epidydimal WAT; BAT = Brown Adipose Tissue

**(C)** Western blot analysis of AR protein expression in various fat pads of 10-week-old control mice. β-actin was used as a loading control. ING = Inguinal WAT; EP = Epidydimal WAT; BAT = Brown Adipose Tissue
**(D)** Schematic representation of *Ucp1*-CreER$^{T2}$ and floxed (L2) exon 1 *Ar* alleles. LoxP sites are shown by arrowheads. Tamoxifen injections were used to induce CreER$^{T2}$ activity and excise exon 1 (L-). Primers used for allele characterization are depicted in Supplementary Table 1.
**(E)** Representative electrophoresis of PCR-amplified genomic DNA from control and AR$^{iB/-y}$ mice. Distinct amplicon sizes correspond to AR L2 (250 bp) and AR L- (500 bp).
**(F-G)** Hematoxylin and Eosin (H&E) staining of epWAT **(F)** and BAT **(G)** from control and AR$^{iB-/y}$ mice at 9 and 10 weeks of age. Insets at the lower-left corner represent a higher magnification of corresponding images. Scale bars, 100 µm.
**(H-I)** Quantification of body **(H)** and fat pad weights (BAT, epWAT, ingWAT) **(I)** at 10, 15, and 30 weeks of age, for control and AR$^{iB-/y}$ mice. Data are presented as mean + SEM. Statistical test used was 2-way ANOVA with Fischer's LSD test. *, $p < 0.05$; **, $p < 0.01$

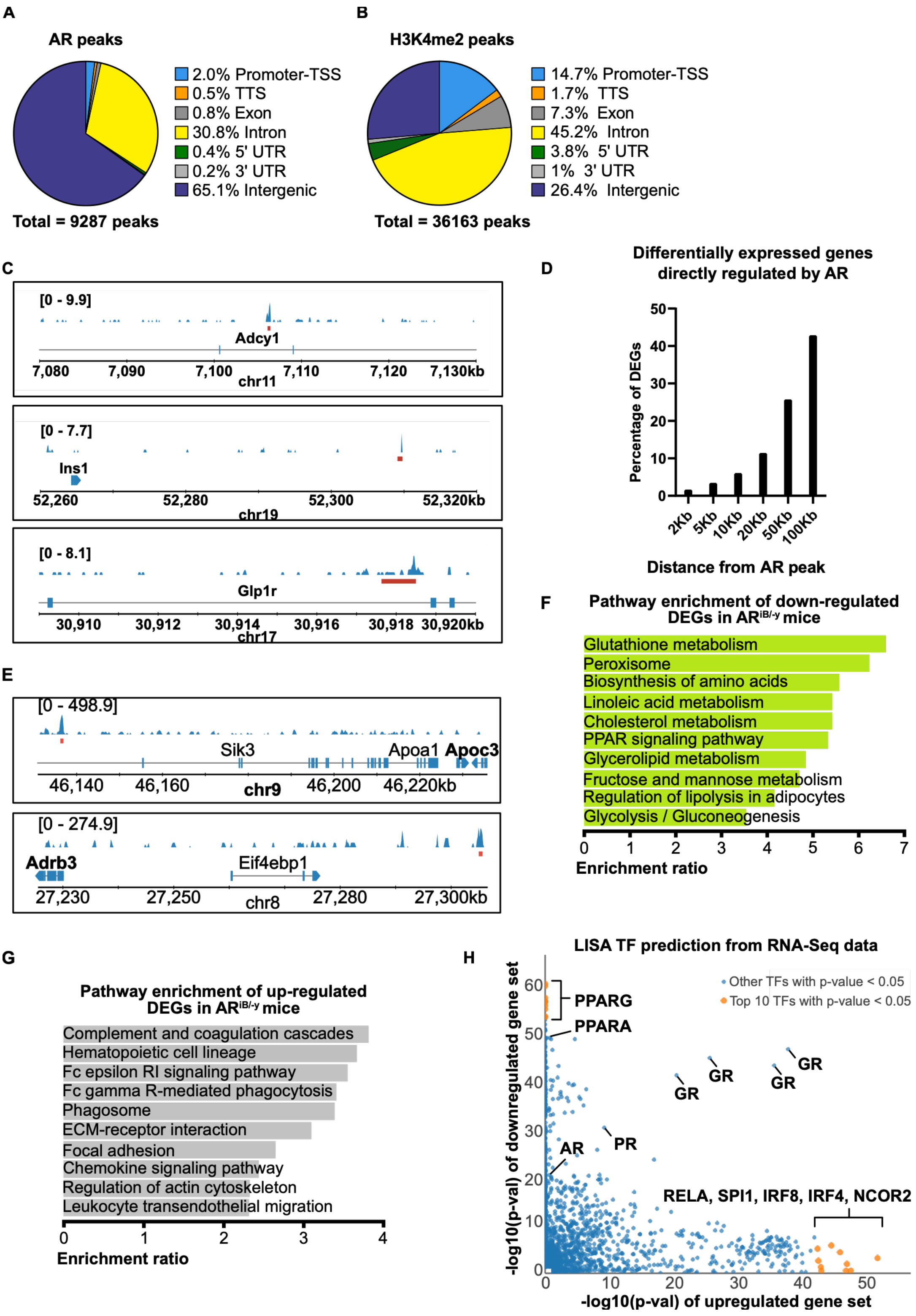


**Figure S2: Characterization of AR cistrome and transcriptome in mouse ingWAT.**

**(A-B)** Pie chart depicting the genomic location of AR (**A**) and H3K4me2 (**B**) in 10-week-old ingWAT of wild-type mice.

**(C)** Localization of AR at representative loci from in ingWAT.

**(D)** Histogram depicting the percentage of differentially expressed genes obtained from RNA-seq dataset of 10-week-old control and AR$^{iB-/y}$ mice, in relation to the proximity of their TSS from the nearest detected AR peak from the ChIP-Seq data of 10-week-old ingWAT in wild-type mice.

**(E)** Localization of AR at representative loci from in ingWAT.

**(F-G)** KEGG pathway enrichment analysis of down-regulated **(F)** and up-regulated **(G)** genes in ingWAT of 10-week-old $AR^{iB-/y}$ mice relative to control mice.

**(H)** LISA transcription factor prediction for potential transcriptional regulators of down-regulated and up-regulated genes in ingWAT of 10-week-old $AR^{iB-/y}$ mice relative to control mice. Selected transcription factors are annotated. Top 10 transcription factors with p-value < 0.05 on either axis are indicated in orange, while the other ones are indicated in blue

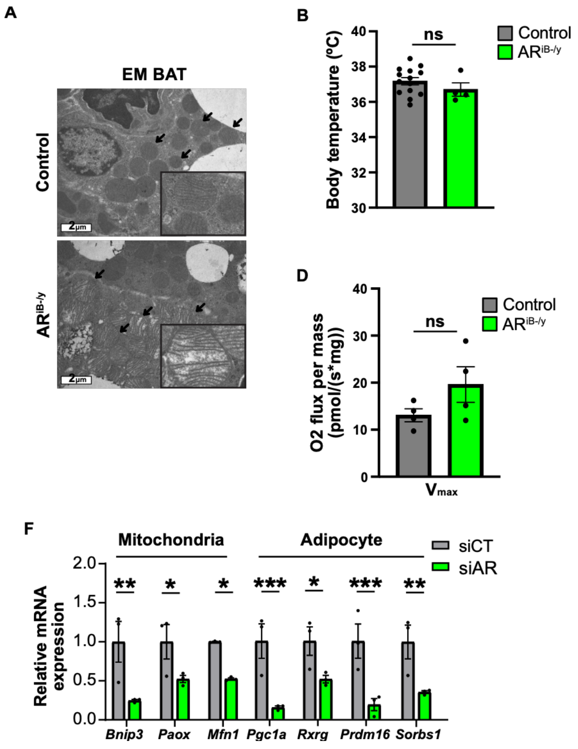


**Figure S3: Consequences of AR ablation in beige adipocytes on mitochondria homeostasis.**

**(A)** Ultrastructure analysis of BAT from 10-week-old control and AR$^{iB-/y}$ mice. Black arrows indicate mitochondria. Insets at the lower-right corner represent higher magnification of corresponding images. Scale bars, 1 µm

**(B)** Quantification of body temperature of 10-week-old control and AR$^{iB-/y}$ mice. Data are presented as mean ± SEM. Statistical test used was two-tailed unpaired t-test. ns: non-significant.

**(C-E)** Quantification of mitochondrial oxygen ($O_2$) flux per mass **(C)**, corresponding maximal respiration **(D)** and relative maximal hydrogen peroxide ($H_2O_2$) flux per unit $O_2$ flux **(E)** under successive supplementation of substrates for different mitochondrial complexes followed by inhibition of complex-I via Rotenone (Rot), in the ingWAT of 10-week-old control and AR$^{iB-/y}$ mice under ambient temperature conditions. Data are presented as mean ± SEM. Statistical test used was multiple two-tailed unpaired t-tests. ns: non-significant. Pyr + Mal = Pyruvate + Malate; ADP = Adenosine Diphosphate; Succ = Succinate.

**(F)** Relative transcript levels for indicated genes in control (siCT) and AR-siRNA transfected (siAR) immortalized beige adipocytes, 3-days post-differentiation. Data are presented as mean ± SEM. Statistical test used was multiple two-tailed unpaired t-tests. *, $p < 0.05$; **, $p < 0.01$; ***, $p<0.001$

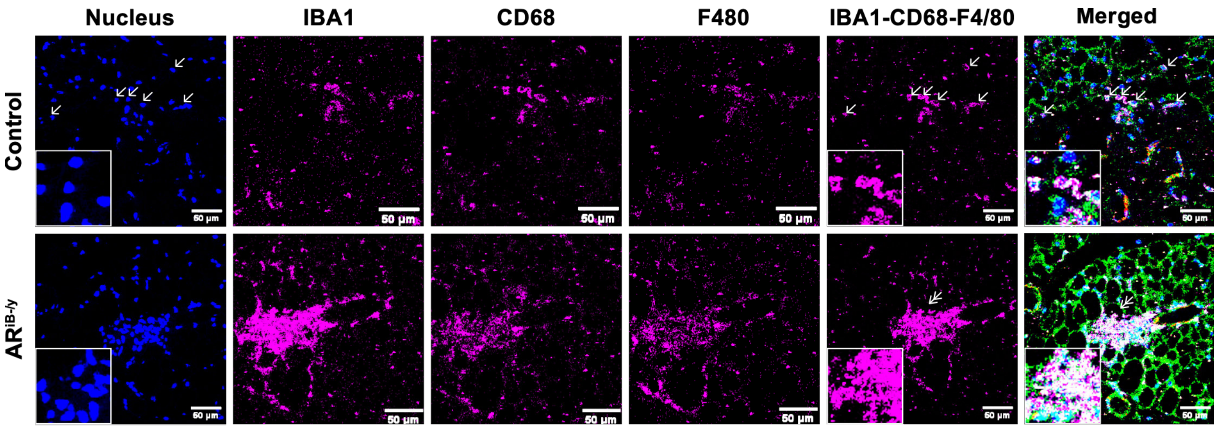


**Figure S4: Characterization of AR invalidation in beige adipocytes on the immune infiltrate.**
Imaging Mass Cytometry detection of IBA1 (magenta), CD68 (magenta), F4/80 (magenta), combination of IBA1-CD68-F4/80 (magenta) and the merged imaged referenced from Figure 4D in ingWAT of 10-week-old control and $AR^{iB-/y}$ mice. Nuclei are visible in blue. Insets at the lower-left corner represent higher magnification of corresponding images. Scale bars, 50 µm.

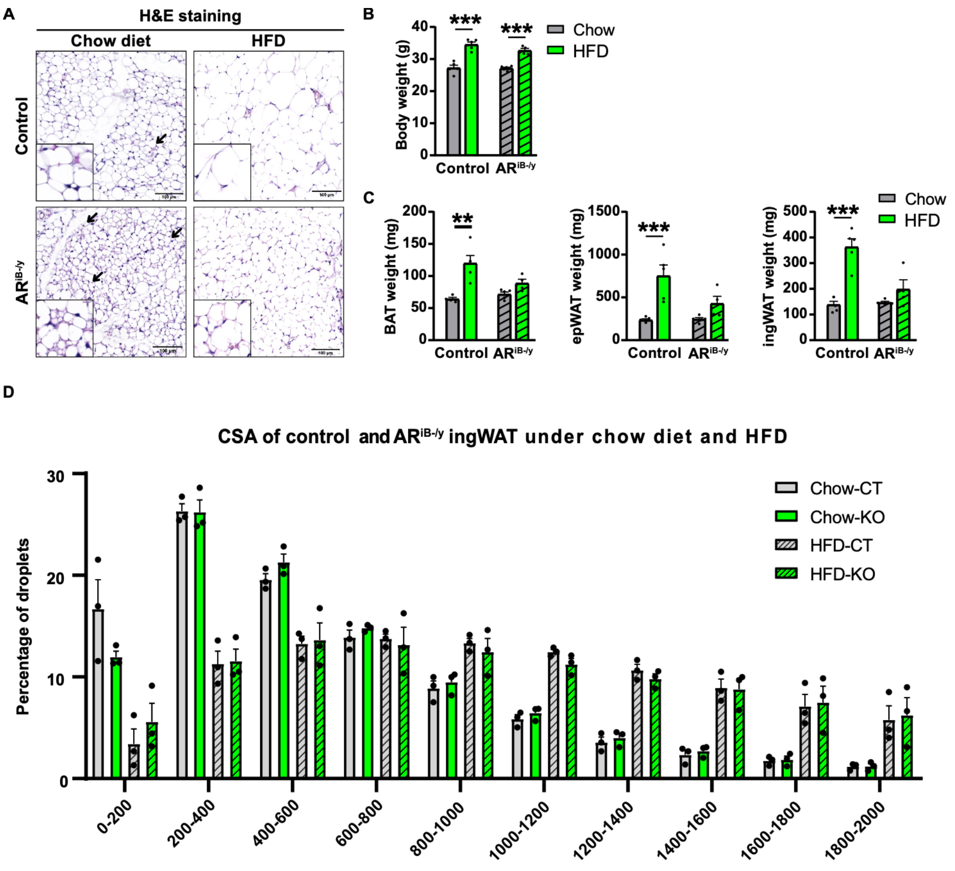


**Figure S5: Impact of AR on the metabolic adaptations following a high-fat diet feeding**

**(A)** Hematoxylin and Eosin (H&E) staining of ingWAT from control and AR$^{iB-/y}$ mice fed a regular chow or a high-fat diet (HFD). Black arrows indicate dense lipid droplet clusters, representative of beige adipocytes. Insets at the lower-left corner represent higher magnification of corresponding images. Scale bars, 100 µm

**(B-C)** Quantification of body **(B)** and fat pad weights (BAT, epWAT, ingWAT) **(C)** of control and AR$^{iB-/y}$ mice fed a regular chow or a HFD. Data are presented as mean + SEM. Statistical test used was 2-way ANOVA with Fischer's LSD test. **, $p < 0.01$; ***, $p<0.001$

**(D)** Quantification of lipid droplet cross-sectional area (CSA) in µm$^2$ in ingWAT of control and AR$^{iB-/y}$ mice under chow diet and HFD. Data are presented as mean + SEM. Statistical test used was two-tailed multiple unpaired t-tests.

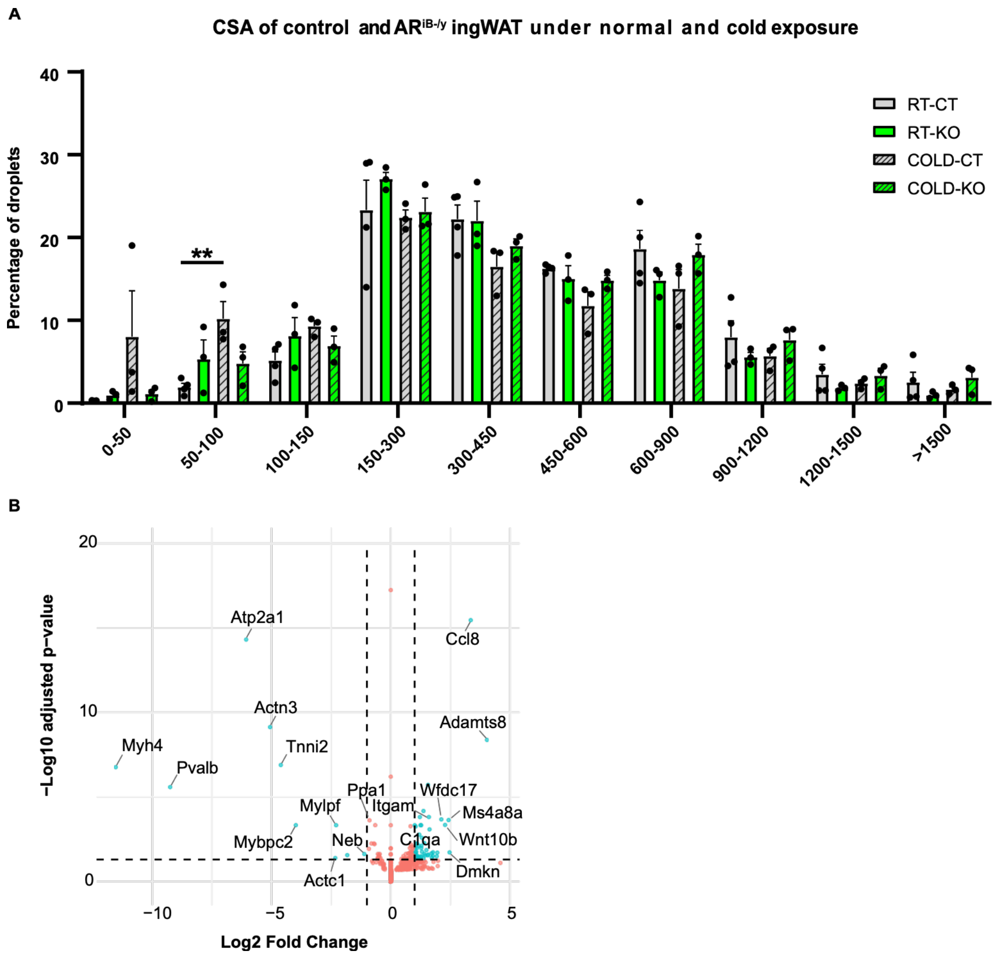


**Figure S6: Impact of AR on the metabolic adaptations following cold exposure**

**(A)** Quantification of lipid droplet cross-sectional area (CSA) in $\mu m^2$ in ingWAT of control and $AR^{iB-/y}$ mice maintained at room temperature (RT) or at 10°C for 10 days (COLD). Data are presented as mean + SEM. Statistical test used was two-tailed multiple unpaired t-tests. **, p<0.01.

**(B)** Volcano plot depicting -log(p-adj) in x-axis and $\log_2$(fold change) of differentially expressed genes from RNA-seq data of ingWAT in cold exposed control and $AR^{iB-/y}$ mice. Selected markers are annotated. Genes with p-adj < 0.05 and | $\log_2$(fold change) | > 1 are indicated in cyan, while the others are indicated in red.

**Supplementary tables**

**Supplementary table 1: List of primers used for genotyping**

| Gene | Forward (5'-3') | Reverse primer (5'-3') |
|---|---|---|
| $CreER^{T2}$ | TTCCCGCAGAACCTGAAGATGTTCG | GGGTGTTATAAGCAATCCCCAGAAATGC |
| *ARL2/WT* | CTGGTTGCTAAGGGACTTCGG | GCCCACACAAACAGTCAGCCCA |
| *ARL-* | GCCCACACAAACAGTCAGCCCA | GCCCACACAAACAGTCAGCCCA |

**Supplementary table 2: List of mouse primers used for RT-qPCR analyses**

| Gene | Forward (5'-3') | Reverse primer (5'-3') |
|---|---|---|
| *18S* | TCGTCTTCGAAACTCCGACT | CGCGGTTCTATTTTGTTGGT |
| *Bnip3* | TTCCACTAGCACCTTCTGATGA | GAACACCGCATTTACAGAACAA |
| *Paox* | AGTCTTCACATGTGCTCTGTGGGT | TGGCAATTGTGGGTTTCCTGTCAC |
| *Mfn1* | CCTCCATGGGCATCATCGTT | TGCAGCTTCTCGGTTGCATA |
| *Ppargc1a* | AAGTGTGGAACTCTCTGGAACTG | GGGTTATCTTGGTTGGCTTTATG |
| *Rxrg* | GGAGACTCTTCGAGAGAAGGTTTAT | ATGAGCTTGAAGAAGAAGAGGTGT |
| *Prdm16* | TGTCAAGGTGTTCACGGACC | GCTGTTTGAGGCCAGAGGAT |
| *Sorbs1* | TACCGAGCGATCGAAAGACT | AGGAATATCGAGGGGAATGG |

**Supplementary table 3: List of antibodies and their metal conjugates used in imaging mass cytometry**

| Metal Tag | Target |
|---|---|
| 111 Cd | COX4I2 |
| 112 Cd | HK2 |
| 113 Cd | 5hmC |
| 115 In | CS |
| 141 Pr | IBA1 |
| 142 Nd | H3K9me3 |
| 143 Nd | GFAP |
| 144 Nd | CD36 |
| 145 Nd | 4EBP1 |
| 146 Nd | BCAT2 |
| 147 Sm | GLUT5 |
| 148 Nd | H3K9ac |
| 149 Sm | H3K9me1 |
| 150 Nd | SDHA |
| 151 Eu | CD31 |
| 152 Sm | pAKT |
| 153 Eu | VDAC1 |
| 154 Sm | OLIG2 |
| 155 Gd | ACADM |
| 156 Gd | LDHA |
| 158 Gd | ECADHERIN |
| 159 Tb | CD68 |
| 160 Gd | ATP5A |
| 161 Dy | HK1 |
| 162 Dy | G6PD |
| 163 Dy | PDHA1 |
| 164 Dy | OPA1 |
| 165 Ho | GLUT1 |
| 166 Er | GPT2 |
| 167 Er | PC |
| 168 Er | KI67 |
| 169 Tm | PKM2 |
| 170 Er | F480 |
| 172 Yb | CASPASE3 |
| 173 Yb | GOT2 |
| 175 Lu | GLUTAMINE SYNTHETASE |
| 176 Yb | CPT1a |
| 191 Ir | DNA1 |
| 193 Ir | DNA2 |
